# White adipose tissue browning preserves UCP1-dependent nonshivering thermogenesis upon brown-adipocyte specific lipase-deficiency

**DOI:** 10.1101/2024.12.04.626093

**Authors:** Yelina Manandhar, Anita Pirchheim, Peter Hofer, Wolfgang Krispel, Nemanja Vujic, Dagmar Kolb, Martina Schweiger, Ulrike Taschler, Robert Zimmermann, Gerald Hoefler, Dagmar Kratky, Rudolf Zechner, Renate Schreiber

## Abstract

Intracellular fatty acids (FAs) activate and fuel non-shivering thermogenesis (NST) via uncoupling protein 1 (UCP1). Adipose triglyceride lipase (ATGL) and hormone-sensitive lipase (HSL) control FA availability. Since mice lacking ATGL in brown adipose tissue (BAT) exhibit intact recruitable adrenergic thermogenesis, we hypothesized that HSL-mediated FA release is sufficient to activate UCP1-dependent NST. We demonstrate that mice with inducible brown adipocyte-specific loss of ATGL and HSL (iBDKO) exhibit normal recruitable adrenergic thermogenesis upon prolonged cold exposure. Mechanistically, we show that BAT thermogenic capacity is impaired in cold-adapted iBDKO mice due to diminished mitochondrial numbers. Increased browning of white adipose tissue (WAT) in iBDKO mice indicates a shift in thermogenesis from BAT to WAT. Consistently, the loss of ATGL and HSL in BAT and WAT disrupts thermogenesis in both depots, resulting in blunted UCP1-dependent NST. Our study highlights the metabolic adaptability of adipose tissue and the critical role of intracellular lipolysis in regulating thermogenesis.

## Introduction

Homeotherms maintain their body temperature within a narrow range despite varying ambient temperatures. Heat production can occur by several means, depending on the length of the cold stimulus. Initially, upon cold exposure for a few hours to days (acute cold), the muscle produces heat via ATP-dependent shivering thermogenesis. When exposed to cold temperature for several weeks (cold adaptation), mammals generate heat via non-shivering thermogenesis (NST), whereby chemical energy is dissipated as heat. The main form of NST is mediated by uncoupling protein 1 (UCP1), which facilitates mitochondrial respiration without ATP synthesis. This UCP1-dependent NST – often also referred to as classical adaptive NST – occurs in brown and beige adipocytes within inguinal white adipose tissue (WAT). UCP1-dependent NST is fueled by circulating glucose, fatty acids (FAs), and triglyceride-rich lipoproteins (TRLs)^1^. Thus, increased adipocyte thermogenesis is associated with improved glucose and lipid control, as well as improved cardiometabolic health^2,3^.

Despite being identified and purified in the early 1980s^4^, the mechanism of action of UCP1 is still not fully understood^5–8^. FAs play a dual role in UCP1-dependent NST as they i) fuel the electron transport chain and ii) activate UCP1^5,6,9–11^. Thereby, UCP1 likely acts as a FA-anion/H^+^ symporter^5^ triggering proton transport via the inner mitochondrial membrane to release energy as heat. It is thought that the FAs originate from the degradation of intracellular triacylglycerols (TGs)^12^ via adipose triglyceride lipase (ATGL) and hormone-sensitive lipase (HSL)^13^ in a process termed lipolysis. ATGL and HSL are directly or indirectly controlled by the sympathetic nervous system. Upon sensing cold, terminal axons of sympathetic nerves release catecholamines, which bind to β3-adrenergic receptors in brown adipocytes and initiate downstream signaling pathways to increase cyclic AMP (cAMP) levels and activate protein kinase A (PKA). PKA, in turn, triggers several critical phosphorylation and translocation events involving proteins that are essential for lipolysis^14–16^, thereby increasing FA release from TGs stored in cytoplasmic lipid droplets (LDs). PKA also controls the transcription of *Ucp1* and of genes involved in adipocyte differentiation and mitochondrial biogenesis, such as *Pgc1α* and *Ppparγ*, via p38/MAPK^17^ and the cAMP response element (CREB)^18,19^. Together, this coordinated response increases thermogenic capacity in brown adipocytes.

Studies in mice lacking ATGL-mediated lipolysis in brown adipose tissue (BAT) challenged the dogma that FAs derived from intracellular lipolysis are essential to activate UCP1-dependent NST^20,21^. In the absence of ATGL or its coactivator *α*/β hydrolase domain containing protein 5 (ABHD5, also known as CGI-58) in BAT, mice maintained euthermia in the cold and had normal recruitable adrenergic thermogenesis indicative of UCP1-dependent NST. These findings raised the question of the source of FAs that trigger UCP1-mediated proton transport^22^. Several scenarios are plausible: FAs may derive from lipolysis in WAT or from TRLs to activate and fuel UCP1 in BAT. Alternatively, although exhibiting a weak TG hydrolase activity^23^, HSL may release sufficient FAs within BAT to activate UCP1, for which the FA concentration threshold remains unknown. Yet, HSL cannot compensate for the loss of ATGL in WAT or the heart, i.e. to provide sufficient fuel during fasting^20,24^ or for the activation of nuclear receptors^25^.

Here, we demonstrate that in the absence of both ATGL and HSL in BAT (tamoxifen-inducible <u>B</u>rown-adipocyte specific <u>D</u>ouble ATGL and HSL <u>K</u>nock-<u>O</u>ut, iBDKO), recruitable adrenergic thermogenesis upon cold adaptation remains intact via a BAT-independent mechanism. The genetic loss of ATGL and HSL reduces the thermogenic capacity of BAT specifically upon cold adaptation due to low mitochondrial content despite an increased number of brown adipocytes. Normal UCP1-dependent NST in iBDKO mice is enabled by an increased thermogenic capacity of inguinal WAT. This rescue mechanism of WAT thermogenesis - when thermogenic capacity in BAT is impaired - is absent when ATGL and HSL are genetically deleted in both BAT and WAT (<u>A</u>dipocyte-specific <u>D</u>ouble ATGL and HSL <u>K</u>nock-<u>O</u>ut, ADKO), thus disabling UCP1-dependent NST. Nevertheless, ADKO mice maintained euthermia in the cold via a currently unknown mechanism. Moreover, we show that iBDKO mice are susceptible to hypothermia when exposed to cold and fasted simultaneously, exclusively upon short-term but not prolonged deletion of ATGL and HSL. Finally, and in contrast to the heart or WAT, our data suggest a functional redundancy of ATGL and HSL in BAT as single lipase deficient mice neither show a BAT defect nor fasting-induced cold sensitivity. Our study highlights the metabolic adaptability of adipose tissue and the critical role of intracellular lipolysis in regulating thermogenesis.

## Results

### Validation of tamoxifen-inducible brown-adipocyte specific double ATGL and HSL knockout (iBDKO) mice

To study the loss of the two major adipocyte TG lipases ATGL and HSL on UCP1-mediated NST, we generated and characterized *Pnpla2*^flx/flx^*Lipe*^flx/flx^::*Ucp1*-CreER^+/-^ mice (tamoxifen-inducible <u>B</u>rown adipocyte-specific <u>D</u>ouble ATGL and HSL <u>K</u>nock-<u>O</u>ut mice, synonym: iBDKO; Fig. 1a). Immunoblot analysis of whole BAT homogenates showed reduced protein levels of ATGL (-70%) and HSL (-84%) in iBDKO compared to *Pnpla2*^flx/flx^*Lipe*^flx/flx^ littermate controls (synonym: controls, Fig. 1b). Enzyme activity assays in low-fat infranatants from BAT homogenates showed 60% lower TG hydrolase activities in iBDKO mice than in controls (Fig. 1c). Addition of specific small molecule inhibitors for ATGL (ATGLi) and HSL (HSLi) or the non-specific serine hydrolase inhibitor orlistat reduced TG hydrolase activities in BAT samples from controls by 85% and 91%, respectively. The addition of these inhibitors to BAT samples from iBDKO reduced TG hydrolase activities to the levels observed in control samples. These results suggest that there is residual active ATGL and HSL protein in bulk BAT homogenates of iBDKO mice. Morphologically, BAT of iBDKO mice appeared whitened (Fig. 1d, upper panel) with a ∼5-fold increase in BAT mass (Fig. 1e). Histological analysis revealed large and unilocular LDs throughout the BAT sections of iBDKO mice compared to the characteristic multilocular LDs in the BAT of the controls (Fig. 1d, lower panel). Taken together, these data suggest that the residual lipase protein expression and lipolytic activities in bulk BAT homogenates of iBDKO mice is likely derived from WAT surrounding the BAT and/or from non-adipocyte cells within BAT.

**Fig. 1.**
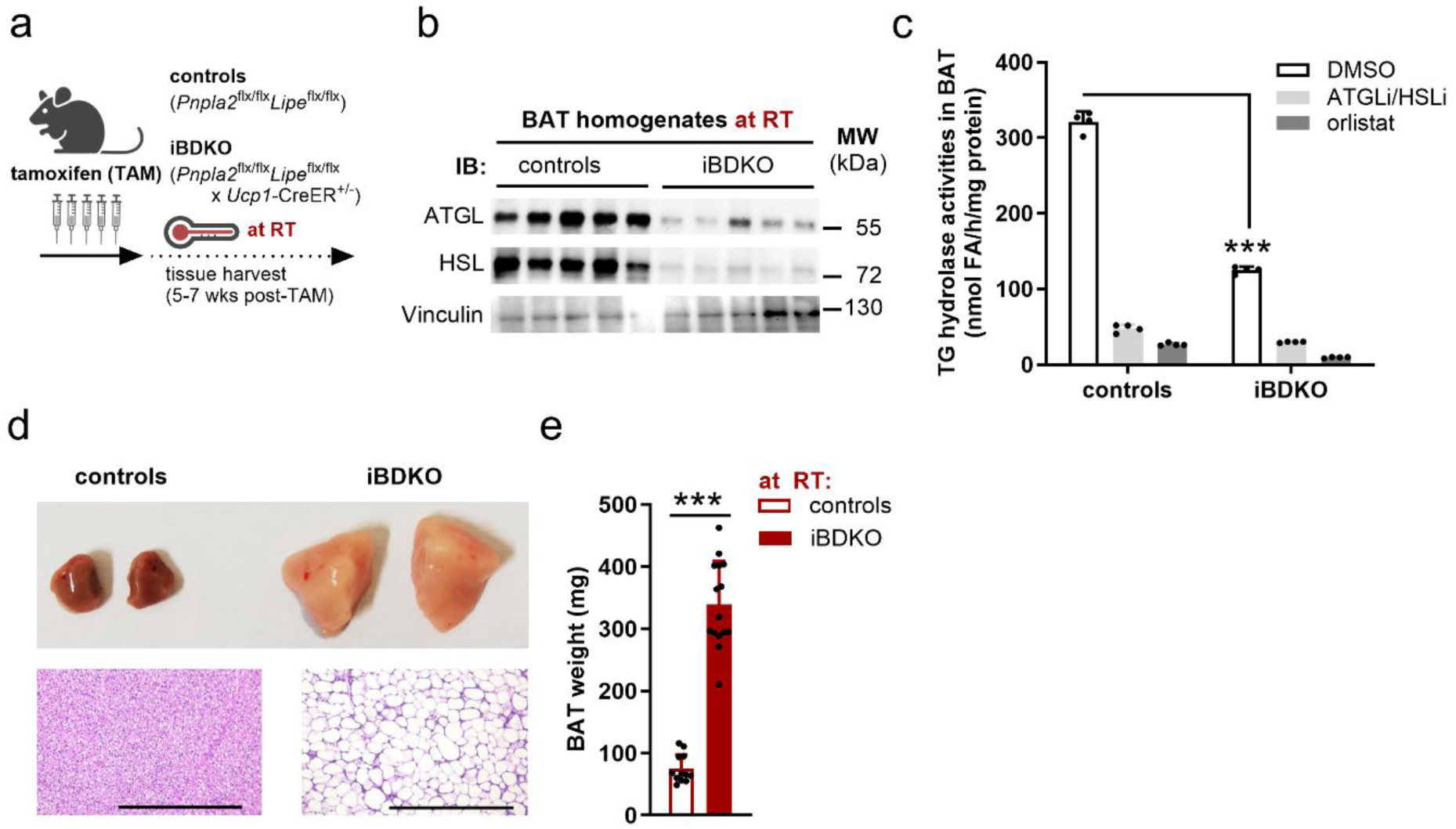
Genetic loss of ATGL and HSL in brown adipocytes causes BAT hypertrophy. Tamoxifen-inducible <u>B</u>rown-adipocyte specific <u>D</u>ouble ATGL and HSL <u>K</u>nock-<u>O</u>ut mice (iBDKO) were generated by crossing *Pnpla2*^flx/flx^*Lipe*^flx/flx^ with *Ucp1*-CreER^+/-^ mice. Gene deletion was induced in adult male mice by intragastric gavage of tamoxifen (TAM) for 5 subsequent days. (**a**) Schematic presentation of the mouse model studied and the tamoxifen treatment. (**b**) Immunoblot (IB) analyses from total BAT homogenates. Delipidated volume equivalents of 0.25% from total BAT were loaded per lane. (**c**) TG hydrolase activities were determined in a pooled fraction of fat-free infranatants from BAT homogenates (n = 7) using a radiolabeled triolein substrate emulsified with phospholipids. Enzymatic activities were determined in the absence or presence of inhibitors specific for ATGL (ATGLi) or HSL (HSLi) or the non-specific serine hydrolase inhibitor orlistat. Technical replicates are shown. (**d**) Gross morphology and histology (H/E stain) of BAT (upper and lower graph, respectively). Bar graph is 500 µm. (**e**) BAT weights. n = 12-15. Data are means + SD from male mice housed at ambient room temperature of 21-23 °C and 6-14 weeks post-TAM. Statistical analyses were performed using Student’s *t*-test. ***, *p*<0.001.

### iBDKO mice show increased body temperature and intact recruitable adrenergic thermogenesis upon cold adaptation

To assess the effect of inducible brown adipocyte-specific ATGL and HSL deficiency on body temperature during cold exposure at 5 °C, we implanted telemetry devices to measure core body temperature. A scheme illustrating the experimental settings is shown in Fig. 2a. iBDKO mice fed *ad libitum* tolerated cold exposure well (Fig. 2b). In fact, upon prolonged cold exposure, iBDKO core body temperature was +0.8 °C and +0.3 °C higher than in controls during day and night, respectively (Fig. 2b, right graph). The higher core body temperature in *ad libitum* fed iBDKO mice adapted within the first two days of cold exposure (Fig. 2b, left graph). At room temperature (RT), core body temperature in iBDKO was +0.1 °C higher during the day and unchanged during the night (Extended Data Fig. 1a).

**Fig. 2.**
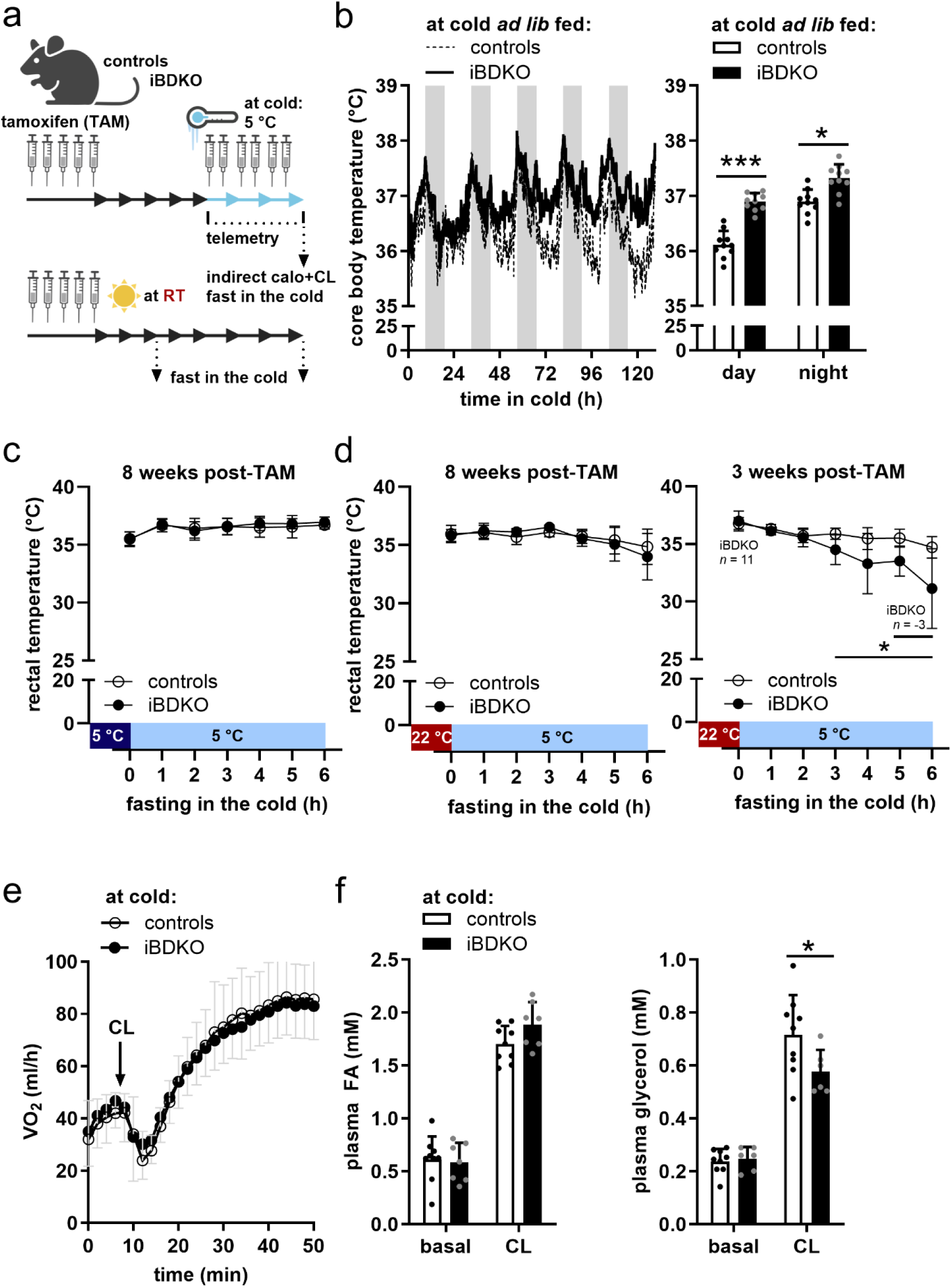
iBDKO mice exhibit increased core body temperature and intact recruitable adrenergic thermogenesis i*n vivo* upon cold adaptation. (a) Schematic presentation of the mouse model studied, tamoxifen treatment, and experimental settings. Blue thermometer indicates cold exposure at 5 °C. Arrows indicate weeks. Dashed lines (±arrows) indicate time-point/-period of experiments. (**b**) Time-course (left) and mean (right) of core body temperature at 5 °C (n = 10). Non-shaded and grey shaded areas represent day and night, respectively. (**c**) Body temperature from mice previously housed at 5 °C for ≥3 weeks (dark blue area) and exposed to 5 °C during fasting (light blue area, n = 10-11). (**d**) Body temperature from mice 8 weeks (left graph) or 3 weeks (right graph) after gene deletion using tamoxifen (post-TAM). Mice were previously housed at 22-23 °C (red area) and acutely shifted to 5 °C during fasting (light blue area, n = 10-11). (**f**) Oxygen consumption (VO_2_) during unstimulated conditions and upon CL316,243 (CL) injection in pentobarbital-anesthetized mice upon cold adaptation (n = 12-13). VO_2_ was analyzed at an ambient temperature of 28-30 °C. (**g**) Plasma FA and glycerol levels during unstimulated (basal) conditions and after CL injection (n = 7-9). Mice were implanted with telemetry transmitters to assess core body temperature and exposed to 5 °C for ≥3-weeks. Body temperature was assessed using a rectal probe. Data are means + SD (except for time-course in B showing mean only) from male mice aged 19-28 weeks and 7-11 weeks post-TAM. Statistical analyses were performed using Student’s *t*-test. *, *p*<0.05 and ***, *p*<0.001.

Recently, Mouisel *et al*.^26^ showed severe hypothermia in iBDKO mice upon cold exposure and concomitant fasting. When we repeated this experiment in mice 8 weeks after gene deletion using tamoxifen (post-TAM) and concomitant cold adaption for 3 weeks (dark blue area on x-axis), both control and iBDKO mice maintained euthermia in a cold environment despite food deprivation (Fig. 2c). When mice that were previously housed at RT (red area on x-axis) and shifted acutely to a cold environment with concomitant fasting (Fig. 2d), the ability to maintain body temperature depended on the length of gene deletion. After 8 weeks of TAM, iBDKO mice maintained euthermia (Fig. 2d, left), but upon 3 weeks post-TAM, iBDKO mice developed hypothermia as soon as 4 h of cold and some mice (*n* = 3) had to be removed from cold due to a body temperature ≤30 °C (Fig. 2d, right). These data show that iBDKO mice fail to adapt to high metabolic stress and suggest an impaired thermogenesis due to the short-term loss of both ATGL and HSL to provide sufficient FAs to fuel and/or activate UCP1. Next, we delineated whether this time dependence of gene deletion in iBDKO mice on the susceptibility of hypothermia induced by both cold and fasting depended on the loss of ATGL or HSL. Therefore, we studied *Pnpla2*^flx/flx^::*Ucp1*-CreER^+/-^ (tamoxifen-inducible <u>B</u>rown adipocyte-specific <u>A</u>TGL <u>K</u>nock-<u>O</u>ut; synonym: iBAKO^20^) and the newly generated *Lipe*^flx/flx^::*Ucp1*-CreER^+/-^ (tamoxifen-inducible <u>B</u>rown adipocyte-specific <u>H</u>SL <u>K</u>nock-<u>O</u>ut; synonym: iBHKO) mice as well as their respective floxed littermates (controls). To assess whether the tamoxifen treatment itself – independent of gene deletion – causes cold intolerance, we included *Ucp1*-CreER^+/-^ as additional controls. Similar to iBDKO, these mice were previously housed at RT and shifted acutely to a cold environment at 5° with concomitant fasting. Neither iBAKO, *Ucp1*-CreER^+/-^ (Extended Data Fig. 1b) nor iBHKO (Extended Data Fig. 1c) exhibited cold intolerance upon acute cold exposure and fasting. Taken together, our results show that fasting- and cold-induced hypothermia are specific to a short-term loss of both ATGL and HSL suggesting that intracellular lipolysis is crucial to activate UCP1 in BAT. However, our findings also demonstrate that the maintenance and increase in body temperature in iBDKO mice depended on the length of gene deletion and cold exposure indicative of an adaptive process, such as UCP1-dependent NST.

The sole analysis of body temperature does not allow a conclusion on functional UCP1-dependent NST^19,27^. UCP1-dependent NST is adaptive, i.e., it is activated and new adipocytes differentiate (recruitment) in response to a cold environment for several weeks. Therefore, we first acclimatized mice to cold for ≥3 weeks and then assessed the effect of the β3-adrenergic receptor agonist CL316,243 (CL) on whole-body O_2_ consumption *in vivo*^9,19^. Notably, and similar to observations made in iBAKO mice^20^, whole-body oxygen consumption (VO_2_) was normal in cold-adapted iBDKO mice when compared to controls under both unstimulated (basal) and CL-stimulated conditions (Fig. 2e). Plasma FA and glycerol levels were also comparable between the genotypes (Fig. 2f). These data demonstrate that brown adipocyte-specific lipolysis mediated by ATGL and HSL is not essential to allow recruitable adrenergic thermogenesis *in vivo* indicative of UCP1-dependent NST.

To assess whether the brown adipocyte-specific loss of ATGL and HSL affects the metabolic phenotype, we performed indirect calorimetry at 5 °C in cold-adapted mice. The respiratory exchange ratio (RER) was higher during the day and lower during the night in iBDKO mice than in controls (Extended Data Fig. 1d). This attenuated light-induced switch from carbohydrate to fat oxidation in iBDKO mice indicated reduced metabolic flexibility. Consistent with this observation, food intake was 32% higher in iBDKO mice during the day but unchanged at night (Extended Data Fig. 1e). Linear regression analyses of energy expenditure (EE) *vs* body weight - similar to ANCOVA^28^ - revealed a trend toward higher EE during the day in iBDKO mice but not at night (Extended Data Fig. 1f). Physical activity was unchanged between the groups (Extended Data Fig. 1g). These results show that brown adipocyte-specific loss of ATGL and HSL causes a marked change in energy metabolism towards increased carbohydrate utilization.

### Cold-adapted iBDKO mice have an impaired BAT thermogenic capacity

A normal adrenergic-stimulated whole-body respiration in iBDKO mice argued for an intact UCP1-dependent NST^27^. We therefore assessed whether this *in vivo* analysis is reflected by normal BAT activity. Histology revealed a heterogeneous cellular pattern with brown adipocytes of different LD sizes and various cell types present in BAT sections from iBDKO mice (Fig. 3a, top). Gene expression analyses confirmed a marked increase of fibrosis and inflammation markers in BAT of iBDKO mice. This increase was independent of cold adaptation and also present in mice kept at RT (Extended Data Fig. 2a). Collagen depositions in BAT of iBDKO were visible in transmission electron microscopy micrographs (TEM, Extended Data Fig. 2b). Similar as in RT and in cold-adapted mice, immunoblot analyses confirmed lower ATGL (-90%) and HSL (-85%) protein expression in whole BAT homogenates of iBDKO mice (Extended Data Fig. 2c), confirming maintained gene deletion due to continued tamoxifen treatment.

**Fig. 3.**
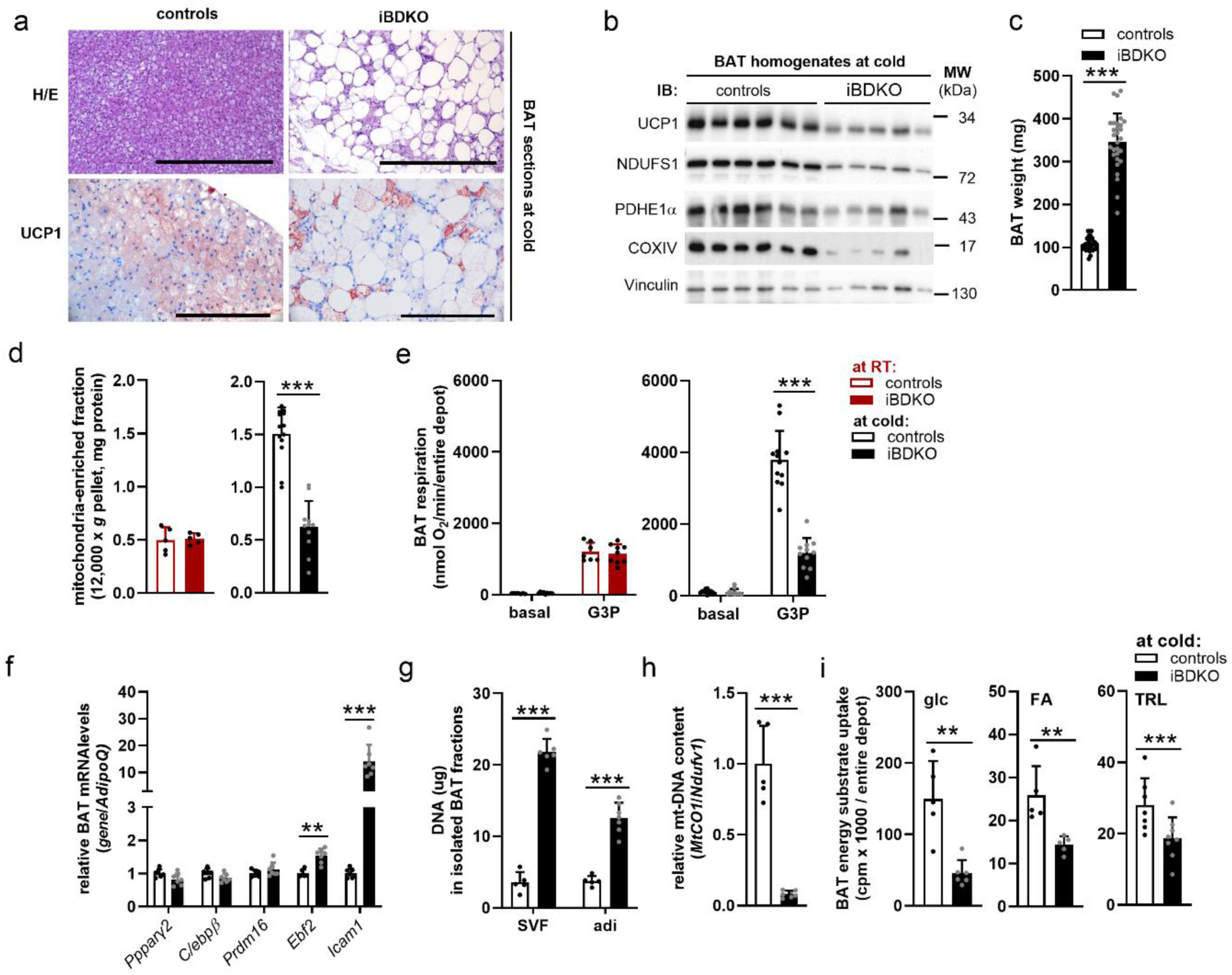
iBDKO mice have impaired BAT thermogenic capacity upon cold adaptation. (**a**) Histology (H/E stain, top; bar graph = 500 µm) and UCP1 immunohistochemistry (bottom; bar graph = 200 µm) in BAT sections of cold-adapted control and iBDKO mice. (**b**) Immunoblot (IB) of BAT homogenates upon cold adaptation. Delipidated volume equivalents of 0.25% of total BAT were loaded per lane. (**c**) BAT weights (n = 29). (**d**) Protein content in the mitochondria-enriched fraction (12,000 x *g* pellet) isolated from BAT of mice housed at room temperature RT (n = 5) and upon cold adaptation (n = 11-13). (**e**) Respiration using glycerol-3-P (G3P) as substrate in BAT homogenates from mice housed at RT (n=7-9) and upon cold adaptation (n = 11-12). (**f**) Gene expression in BAT (n = 6-7). (**g**) DNA content in the isolated stromavascular fractions (SVF) and brown adipocytes (adi) from whole BAT depots of cold-adapted mice (n = 5-7). (**h**) Relative mitochondrial content in isolated brown adipocytes (n = 5-7). (**i**) Energy substrate uptake in BAT using ^3^H-2-deoxyglucose (glc, n = 5-6), ^14^C-*R*-2-bromo palmitic acid (FA, n=5), and ^3^H-TG-rich lipoproteins (TRL, n = 7-8) upon cold adaptation. RT-housed male mice were 17-25 weeks of age and 6-8 weeks post-TAM. Cold-adapted male mice were 16-24 weeks of age and 6-11 weeks post-TAM that were cold-adapted at 5 °C for ≥3 weeks. Data are means + SD. Statistical analyses were performed using Student’s *t*-test. **, *p*<0.01 and ***, *p*<0.001.

The thermogenic capacity of BAT was assessed by whole tissue UCP1 protein content, mitochondrial function, and substrate uptake analyses. Immunohistochemistry showed low overall UCP1 protein expression in BAT sections of iBDKO mice (Fig. 3a, bottom). To quantitate protein expression as indicator of whole BAT thermogenic capacity^29^, tissue volume equivalents were loaded for immunoblot analyses to account for differences in BAT mass and cellular composition between genotypes. The results confirmed lower protein content of UCP1 (-66%) and other mitochondrial markers (≤60%) in BAT of iBDKO mice upon cold adaptation (Fig. 3b), which was not observed when mice were housed at RT (Extended Data Fig. 2d). Upon cold adaptation, BAT mass was ∼3-fold higher in iBDKO mice than in controls (Fig. 3c). As a measure of brown adipocyte differentiation/recruitment, BAT mass increased by ∼50% in control mice upon cold adaptation compared to RT (compare Fig. 1e to Fig. 3c), suggesting the expected brown adipocyte recruitment (105±15 mg at cold *vs* 75±23 mg at RT, *p*<0.001). In contrast, no difference in BAT mass was observed between iBDKO mice housed upon cold adaptation or at RT (339±70 mg at cold *vs* 347±66 mg at RT, *p*=0.7). Similar to mitochondrial protein abundance, total protein content of the mitochondria-enriched fraction from the entire BAT depot was unchanged between genotypes when mice were housed at RT, but 60% lower in iBDKO mice upon cold adaptation (Fig. 3d). Concomitantly, mitochondrial respiration in BAT homogenates using glycerol-3-phosphate (G3P) as substrate was unchanged when mice were housed at RT, but 70% lower in cold-adapted iBDKO mice compared to controls (Fig. 3e). Upon cold adaptation, control mice showed the expected increase in respiration as compared to RT, consistent with brown adipocyte recruitment. In contrast, iBDKO mice showed a similar respiration upon cold adaptation as at RT. Mitochondrial protein expression was not different in isolated mitochondria from BAT upon cold adaptation (Extended Data Fig. 2e). TEM analysis suggested less mitochondria that were squeezed between LDs in BAT of iBDKO, but with normal mitochondrial morphology (Extended Data Fig. 2b). In contrast, none of these differences in mitochondrial markers, content, or respiration were observed in cold-adapted BAT of iBAKO^20^ (Extended Data Fig. 2f) or iBHKO mice (Extended Data Fig. 2g).

The impaired thermogenic capacity in BAT of iBDKO upon cold adaptation may be due to decreased brown adipocyte recruitment. Genes involved in brown adipocyte differentiation were similarly expressed in BAT of iBDKO mice and controls (Fig. 3f). However, the expression of *Icam1* was higher, suggesting a greater number of adipocyte progenitors. We fractionated BAT into the stromavascular fraction (SVF) containing all non-adipocytes, and brown adipocytes (adi) from mice upon cold adaptation and isolated DNA as a measure of cell number. The DNA content in isolated SVF was 6-fold higher in iBDKO (Fig. 3g) than in controls as already indicated by the elevated expression of fibrotic and inflammatory genes (Extended Data Fig. 2a) in BAT of iBDKO mice. The DNA content in the brown adipocyte fraction was 3.4-fold higher in iBDKO mice than in controls, demonstrating brown adipocyte hyperplasia upon BAT-specific loss of ATGL and HSL. Finally, we assessed the relative mitochondrial content in the isolated brown adipocyte fractions showing that iBDKO mice have less than 10% mitochondria of controls (Fig. 3h).

Although not a direct measure of BAT activity, but in line with reduced thermogenic capacity, radioactive tracer studies showed an attenuated uptake of glucose (-70%) and FA (-45%, Fig. 3i). Additionally, whole-particle TRL- or FA-uptake from lipoprotein lipase (LPL)-mediated hydrolysis of TRLs was 35% lower in BAT of iBDKO upon cold adaptation (Fig. 3i). Protein levels of GLUT1 were higher, but those of GLUT4, the predominant glucose transporter in BAT, were lower in plasma membrane fractions from iBDKO BAT (Extended Data Fig. 2h). Similarly, protein levels of the FA transporter CD36 were decreased (Extended Data Fig. 2i). LPL protein levels were higher in BAT of iBDKO mice and transcript levels of *Angptl4* - the LPL inhibitory protein - were 5-fold increased (Extended Data Fig. 2i-j). Reduced CD36 and increased *Angptl4* levels are consistent with the reduced FA uptake detected. In line with reduced BAT glucose uptake in iBDKO mice, expression of genes and proteins involved in *de-novo* lipogenesis was blunted (Extended Data Fig. 2k-l).

Together, our findings demonstrate that iBDKO mice are unable to adapt their BAT thermogenic capacity to this environmental challenge of cold adaptation being unable to increase the mitochondrial content.

### Browning of inguinal WAT (ingWAT) in iBDKO mice after prolonged cold exposure

Despite an impaired BAT thermogenic capacity when adapted to a cold environment, iBDKO mice exhibited an intact recruitable adrenergic thermogenesis. These data prompted us to hypothesize that adaptive processes in WAT metabolism may contribute to sustained NST. Indeed, UCP1 protein expression in ingWAT was 4-fold higher in iBDKO as revealed by immunoblotting and IHC (Fig. 4a-b). Protein levels of tyrosine hydroxylase (TH), the rate-limiting enzyme in catecholamine synthesis, were 3-fold higher in ingWAT of iBDKO mice compared to controls during cold (Fig. 4a), indicative of an increased sympathetic tone. Functionally, mitochondrial respiration was 5-times higher in ingWAT homogenates of iBDKO mice than controls upon cold (Fig. 4c). Inhibiting UCP1 by the addition of GDP resulted in 25% inhibition of respiration in controls and 50% reduction in iBDKO ingWAT, confirming higher UCP1 activity in the latter. Energy substrate uptake was also higher for glucose (5.3-fold), FA (1.6-fold), and TRL (4.6-fold) in ingWAT of iBDKO mice (Fig. 4d).

**Fig. 4.**
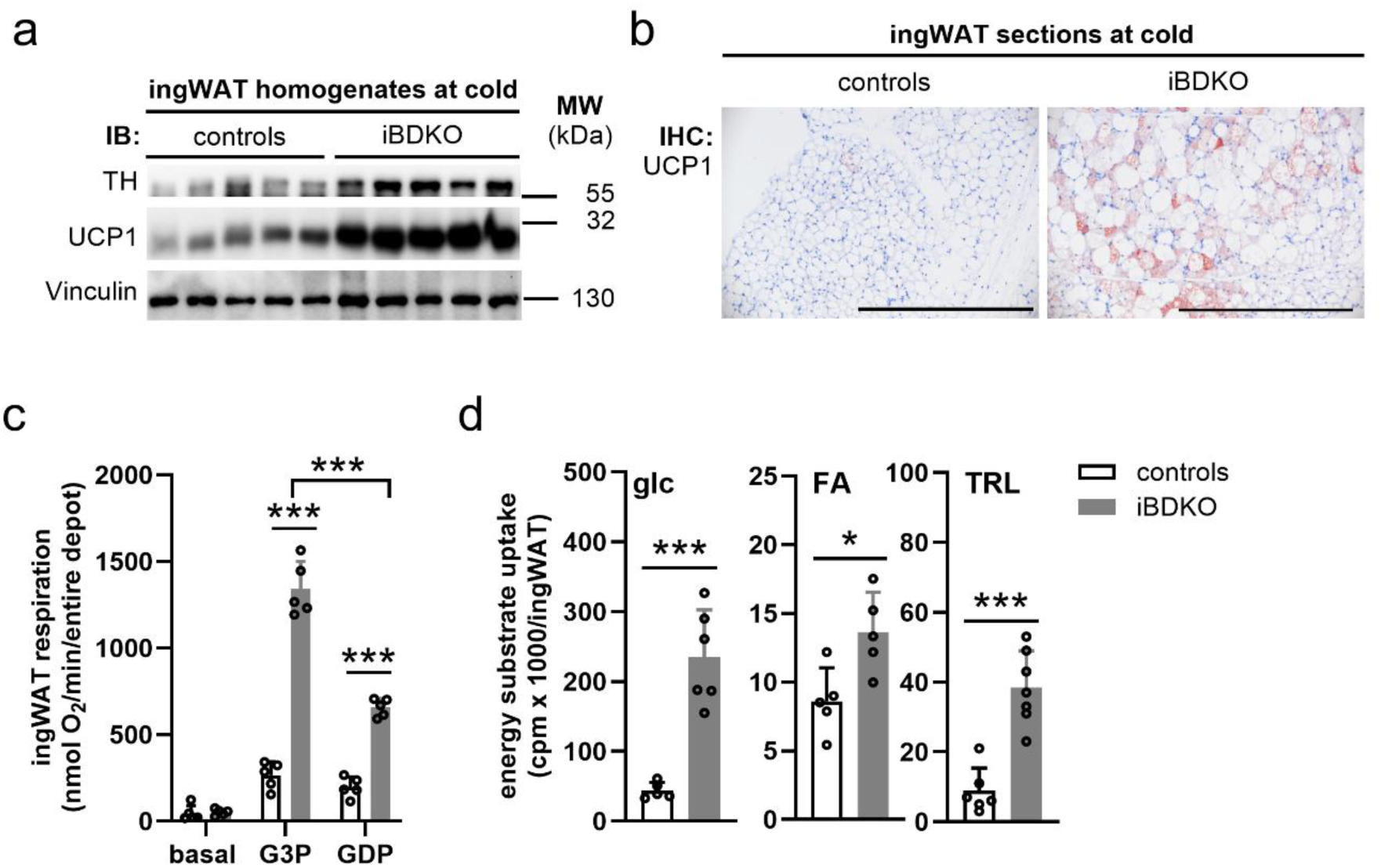
Inguinal white adipose tissue (ingWAT) is browned in iBDKO during cold. (**a**) Immunoblot (IB) of ingWAT homogenates upon cold adaptation. Volume equivalents of 0.25% of total ingWAT were loaded per lane. (**b**) UCP1 immunohistochemistry (IHC) of ingWAT sections. Bar graph = 500 µm. (**c**) Respiration in ingWAT homogenates upon cold adaptation using glycerol-3-P (G3P) as substrate and guanosine-5’-diphosphate (GDP) for UCP1 inhibition (n = 5). (**d**) Energy uptake in ingWAT upon cold adaptation using ^3^H-2-deoxyglucose (glc, n = 6), ^14^C-*R*-2-bromo palmitic acid (FA, n=5), and ^3^H-TG rich lipoproteins (TRL, n = 6-7). Male mice of 16-24 weeks of age and 6-8 weeks post-TAM were exposed to 5 °C for ≥3 weeks. Data are means + SD. Statistical analyses were performed using Student’s *t*-test. *, *p*<0.05 and ***, *p*<0.001.

iBDKO mice harbor an estrogen receptor-dependent Cre-recombinase (Cre) transgene under the transcriptional control of the UCP1 promotor (*Ucp1*-CreER). Increased UCP1 protein expression in ingWAT suggested increased Cre activity in this tissue, which would cause gene and protein deletion of *Pnpla2*/ATGL and *Lipe*/HSL. Indeed, gene expression analyses confirmed that Cre was substantially expressed in ingWAT of iBDKO during cold (Ct values of ∼38 and ∼27 for controls and iBDKO, respectively; Extended Data Fig. 3a). While ATGL mRNA and protein levels were not altered, HSL mRNA and protein levels were decreased by 50% in ingWAT of iBDKO mice (Extended Data Fig. 3a-b). Yet, reduced expression did not translate into reduced lipolytic activities in bulk ingWAT homogenates of iBDKO mice (Extended Data Fig. 3c) possibly because *Ucp1*-Cre is suggested to be less efficient in gene deletion in beige adipocytes^30^.

Collectively, our data suggest that enhanced β3-adrenergic signaling led to increased mitochondrial capacity in ingWAT of iBDKO mice during prolonged cold exposure, which triggered fuel uptake and thermogenesis. Enhanced ingWAT browning may compensate for BAT dysfunction and thus to maintain UCP1-mediated NST in iBDKO mice.

### Genetic loss of ATGL and HSL in white and brown adipocytes (ADKO) impairs UCP1-mediated NST upon cold adaptation

To assess the functional importance of ingWAT browning, we employed a mouse model with white and brown adipocyte-specific loss of ATGL and HSL (*Pnpla2*^flx/flx^*Lipe*^flx/flx^::AdipoQ-Cre^+/-^; <u>A</u>dipocyte-specific <u>D</u>ouble ATGL and HSL <u>K</u>nock-<u>O</u>ut mice, synonym: ADKO; Fig. 5a). These mice lack ATGL and HSL protein as well as TG hydrolase activities in all adipose depots^31^ (Extended Data Fig. 4a-b). Residual TG hydrolase activities in ADKO adipose tissue homogenates were inhibited using the non-specific serine hydrolase inhibitor orlistat (Extended Data Fig. 4b), however *in vivo* lipolysis was not stimulated in ADKO mice compared to controls (Extended Data Fig. 4c), arguing for minor lipase expression in non-adipose cells.

**Fig. 5.**
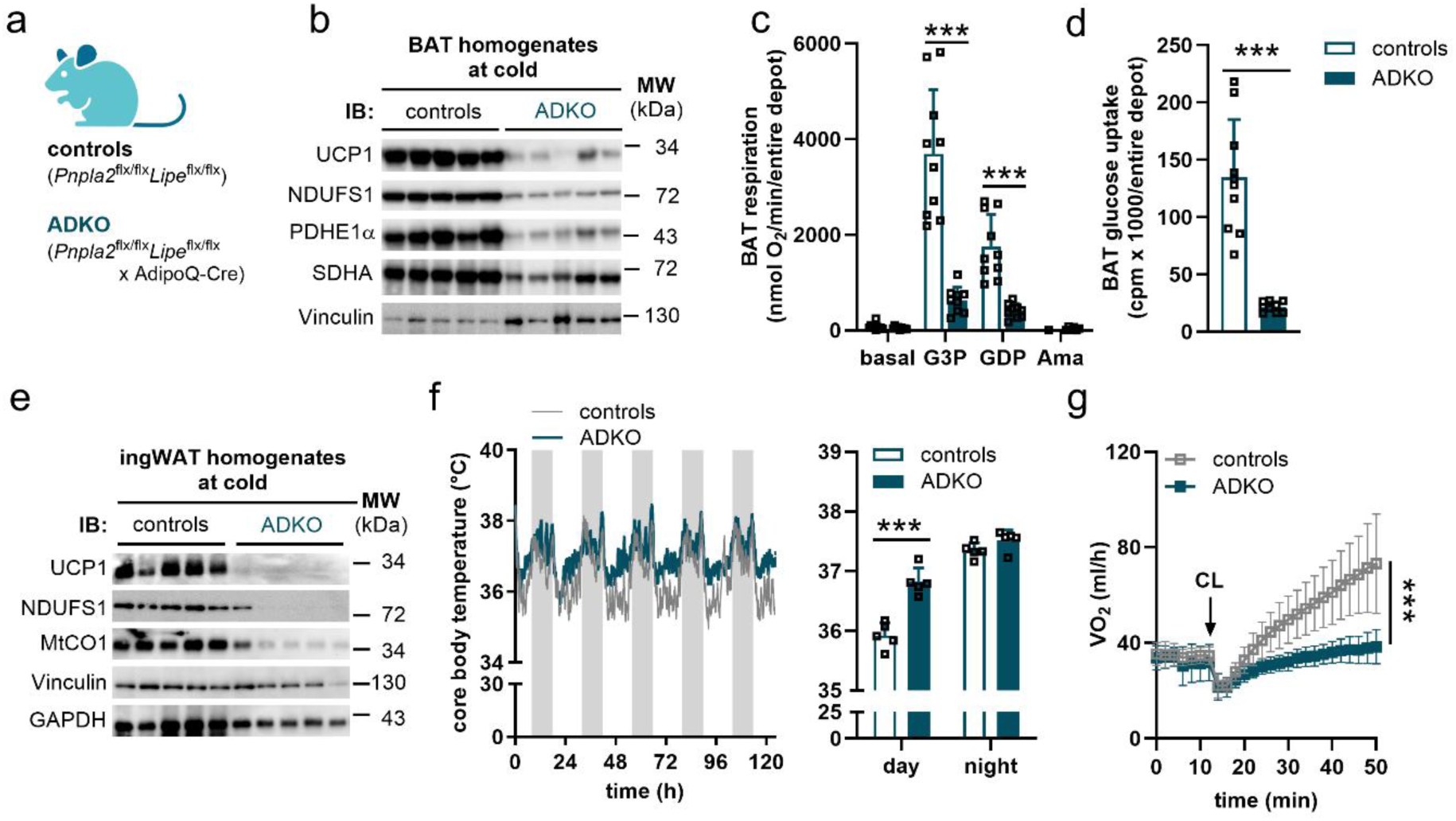
Genetic loss of ATGL and HSL in all adipocytes (ADKO) impairs UCP1-mediated NST in cold. (**a**) Mouse model studied. (**b**) Immunoblot (IB) of BAT homogenates. Volume equivalents of 0.25% of total BAT were loaded per lane. (**c**) Respiration in BAT homogenates (n = 9-10). (**d**) BAT glucose uptake using ^3^H-2-deoxyglucose (n = 9-10). (**e**) Immunoblot (IB) of ingWAT homogenates (5 μg tissue protein per lane). (**f**) Core body temperature using implanted telemetry transmitters. Time-course (left) and mean only (right) at 5 °C (n = 5). Non-shaded and grey shaded areas represent day and night, respectively. (**g**) Oxygen consumption (VO_2_) during unstimulated conditions and upon CL316,243 (CL) injection in pentobarbital-anaesthetized mice upon cold adaptation (n = 7-8). VO_2_ was analyzed at an ambient temperature of 28-30 °C. Male mice aged 14-17 weeks of age and upon cold adaptation at 5 °C for ≥3 weeks were studied. Data are means + SD except otherwise stated. Statistical analyses were performed using Student’s *t*-test. ***, *p*<0.001.

First, we assessed the thermogenic capacity in BAT of ADKO mice by immunoblotting of whole tissue homogenates. ADKO mice exhibited a decrease in protein content of UCP1 (-90%) and other mitochondrial marker proteins (≥70%, Fig. 5b) in BAT homogenates. Concomitantly, BAT respiration and glucose uptake were blunted in cold-adapted ADKO mice by 89% and 84%, respectively (Fig. 5c-d). In contrast to iBDKO, UCP1 and other mitochondrial protein levels (Fig. 5e) as well as respiration (Extended Data Fig. 4d) were reduced by more than 90% in ingWAT of ADKO mice. Accordingly, glucose uptake was 50% lower than in controls (Extended Data Fig. 4e).

To address the impact of the low thermogenic capacity in both BAT and ingWAT on body temperature, we implanted telemetry transmitters and exposed the mice to cold. Remarkably, and similar to iBDKO, ADKO maintained euthermia in the cold and exhibited +0.9 °C higher core body temperature during the day than control mice (Fig. 5f; left, time-course and right, mean). In addition, and similar to iBDKO (Extended Data Fig. 1d), whole-body metabolism was markedly altered in ADKO mice. They showed an attenuated switch of the RER between day and night, indicative of impaired metabolic flexibility (Extended Data Fig. 4f). Food intake was 83% higher during day but unchanged during night (Extended Data Fig. 4g). In contrast to iBDKO mice, energy expenditure was lower during night periods along with attenuated physical activity (Extended Data Fig. 4h-i). Finally, we tested for a recruitable adrenergic thermogenesis as measure for UCP1-mediated NST. In contrast to iBDKO, cold-adapted ADKO mice showed no response to β3-adrenergic receptor activation using CL, suggesting impaired UCP1-mediated NST (Fig. 5g).

Taken together, our data demonstrate that ADKO mice exhibit dysfunctional BAT, and abolished browning of ingWAT, but maintain euthermia when adapted to cold, which is associated with marked whole-body adaptations. Yet, ADKO mice have a blunted recruitable adrenergic thermogenesis indicative of an abolished UCP-1-mediated NST.

## Discussion

FAs have important functions in UCP1-mediated NST, as they fuel the electron transport chain and activate UCP1^6,9–11^. In this regard, FAs derived from intracellular lipolysis have been considered to be essential^19,32,33^. However, we and others showed that UCP1-dependent NST is intact in mice despite absent ATGL-mediated lipolysis in BAT^20,21^. Our studies suggested that WAT- or TRL-derived FAs can sufficiently fuel and activate UCP1. Alternatively, and apart from classical intracellular and vascular lipolysis, lysosomal lipolysis may provide FAs to drive UCP1-dependent NST^34,35^. ATGL and HSL are the two major lipases accounting for more than 90% of TG hydrolase activity in adipocytes^13^ and non-adipocytes. Both lipases can hydrolyze TGs^23^, but ATGL is a more efficient TG hydrolase than HSL^36^. Accordingly, HSL‘s TG hydrolase activity cannot compensate for the loss of ATGL in WAT^20,24,37,38^ or the heart^25^, leading to fasting-induced hypometabolism and cardiomyopathy, respectively. In BAT, it is feasible that HSL activity generates minimal FA levels required to activate UCP1. Here, we assessed how an inducible brown adipocyte-specific loss of ATGL and HSL (iBDKO) affects UCP1-dependent NST and studied these mice upon cold exposure for several weeks (cold adaptation).

We demonstrate that iBDKO mice have an intact UCP1-dependent NST upon cold adaptation. This assumption is based on two key findings: First, within the first days in a cold environment, iBDKO mice develop a higher body temperature than controls, indicative of an evolving adaptive process. And second, after cold adaptation, iBDKO mice exhibit normal recruitable adrenergic thermogenesis. Mechanistically, we show that during cold adaptation the BAT-specific loss of ATGL and HSL blunts the mitochondrial number in BAT leading to a shift in the thermogenic capacity to ingWAT. In the absence of ingWAT browning due to loss of intracellular lipolysis in all fat depots, the recruitable adrenergic thermogenesis *in vivo* is abolished. Moreover, our findings show that iBDKO mice are unable to cope with cold exposure when fasted, leading to hypothermia exclusively upon short-term deletion of both genes. Together, these findings reveal an intact UCP1-mediated NST in mice lacking ATGL and HSL in brown adipocytes, which depends on a thermogenic shift from BAT toward ingWAT.

Our data on intact whole-body respiration in iBDKO mice upon β3-AR stimulation, the key read-out to assess UCP1-mediated NST^27^, contradict a recent publication using the same genetic mouse model^26^. The discrepancies might be explained by different experimental settings: Mouisel *et al*.^26^ studied mice that were housed at RT^26^. We, however, performed the analyses in mice upon cold adaptation to allow the thermogenic capacity to develop^19^. Another mouse model that lacks intracellular lipolysis-derived FAs in BAT are mice with the genetic loss of the two main TG-synthesizing enzymes acyl CoA:diacylglycerol acyltransferases DGAT1 and DGAT2 specifically in brown adipocytes (BA-DGAT-KO model)^39^. These mice exhibited a moderately reduced whole-body response to CL indicative of attenuated UCP1-mediated NST. A reason for the difference between the BA-DGAT-KO and the iBDKO mouse model may be due to germline *vs* inducible gene deletion in adult mice and a lack of compensatory browning of ingWAT in the BA-DGAT-KO model^39^.

Our findings demonstrate that the loss of ATGL and HSL ultimately limited the thermogenic capacity of BAT specifically upon cold adaptation as evident by diminished UCP1 protein levels, oxidative capacity, and fuel uptake. This finding argues that intracellular lipolysis is required to maintain a functional BAT upon high and prolonged metabolic stress. Our data show that brown adipocyte recruitment and mitochondria *per se* are intact in iBDKO mice, but the respiratory capacity is blunted due to a low mitochondrial number in BAT. A reduced mitochondrial content was neither observed by brown adipocyte-specific loss of ATGL alone^20^, HSL alone (this paper), DGAT1/2^39^ nor in whole-body β3-AR deficient mice^40^. All of these models have directly or indirectly impaired intracellular lipolysis in BAT due to the lack of substrate or signaling, respectively. Moreover, our data using metabolic tracers show that iBDKO mice exhibit an impaired BAT uptake of FAs, TRLs and glucose suggesting that FAs derived from these sources as stated by us^20^ or *de novo* lipogenesis^39^ are not an alternative source of FAs for UCP1 activation upon cold adaptation.

Similar to other models of impaired BAT function^41,42^, iBDKO mice showed remarkable browning of ingWAT as evidenced by increased UCP1 protein, mitochondrial respiration, and fuel uptake. Higher TH protein levels indicate an increased sympathetic nerve activity causing the browning effect^43–45^. Indeed, upon chemical denervation of ingWAT^46^ and blocking downstream ATGL-mediated lipolysis^47^, ingWAT browning was absent. In line, ADKO mice lacking ATGL and HSL in both BAT and WAT showed blunted UCP1 and mitochondrial protein levels. This finding raises the question whether ingWAT browning compensates for BAT impairment to maintain normal UCP1-mediated NST in iBDKO mice. Our data in ADKO mice that lack lipolysis in all fat depots support this hypothesis. ADKO mice showed no ingWAT browning associated with blunted UCP1-mediated NST similar as observed in whole-body β3-AR deficient mice^40,48^. These data suggest that ingWAT browning is essential to allow normal UCP1-mediated NST in mice upon loss of ATGL and HSL in BAT. Nevertheless, ADKO mice maintained their body temperature upon cold adaptation, possibly due to adaptive processes like muscle shivering or futile cycles including Ca^2+^ or creatine, while lipid cycling via TG and FAs is impossible due to the lack of lipolysis in all adipocytes. The precise molecular mechanism, however, needs to be determined.

iBDKO mice are susceptible to hypothermia under one specific condition, i.e. when subjected to cold exposure in the absence of food, consistent with the recent findings by Mouisel *et al.*^26^. We show that this cold defect is specific to a short-term loss of 3 weeks of both ATGL and HSL in BAT, but is absent upon single lipase deficiency. These data suggest that i) ATGL and HSL exhibit an apparent functional redundancy in BAT which is in contrast to the heart^25^ or WAT^20,24,37,38^. And ii) FAs derived from ATGL- and HSL-mediated lipolysis are critical to activate UCP1. However, there are some caveats to the later assumption: Compared to fasting, cold exposure was shown to be the superior trigger to increase norepinephrine turnover in BAT^49^, suggesting that cold exposure alone is sufficient to drive intracellular lipolysis. Thus, if FAs derived from ATGL- and HSL-mediated lipolysis are essential to activate UCP1, iBDKO mice would also develop hypothermia upon cold exposure in the fed state, which is not the case^26^.

In contrast to short-term gene deletion, iBDKO mice do not develop hypothermia when subjected to cold exposure and food deprivation simultaneously upon prolonged gene deletion of 8 weeks and independent of the previous housing temperature. These findings suggest an adaptive process in iBDKO mice. In line, neither BA-DGAT-KO mice that lack TGs and thus the substrate for ATGL and HSL in BAT, nor UCP1-deficient mice are susceptible to fasting-induced hypothermia during cold exposure^39,42^. Possible explanations for the latter two mouse models are that alternative mechanisms evolved due to germline deletion like increased *de novo* lipogenesis as an alternative source for FAs to induce UCP1^39^ and UCP1-independent thermogenesis including futile cycles^42,50–52^.

In summary, our data provide evidence that UCP1-mediated NST is intact in the absence of neutral lipolysis in brown adipocytes. The thermogenic capacity is, however, shifted from BAT toward WAT.

## METHODS

### Mouse models

#### BAT-specific knockout models using Ucp1-CreER

Tamoxifen-inducible <u>B</u>rown adipocyte-specific <u>D</u>ouble ATGL and HSL <u>K</u>nock-<u>O</u>ut mice (iBDKO) mice were generated by intercrossing *Pnpla2*^flx/flx^ ::*Ucp1*-CreER^+/-^ (iBAKO^20^) with *Lipe*^flx/flx^ mice^53^ (The Jackson Laboratory stock number 019004) to obtain *Pnpla2*^flx/flx^*Lipe*^flx/flx^::*Ucp1*-CreER^+/-^. *Pnpla2*^flx/flx^*Lipe*^flx/flx^ littermates were used as controls. Tamoxifen-inducible <u>B</u>AT-specific single <u>A</u>TGL <u>K</u>nock-<u>O</u>ut mice (iBAKO; *Pnpla2*^flx/flx^::*Ucp1*-CreER^+/-^) mice were generated as described^20^. *Pnpla2*^flx/flx^ (The Jackson Laboratory stock number 024278) littermates were used as controls. Tamoxifen-inducible <u>B</u>rown adipocyte-specific <u>H</u>SL <u>K</u>nock-<u>O</u>ut mice (iBHKO) mice were generated by intercrossing *Lipe*^flx/flx^ mice with *Ucp1*-CreER^+/-^ to obtain *Lipe*^flx/flx^::*Ucp1*-CreER^+/-^.

#### Adipocyte-specific knockout models using Adiponectin-Cre

<u>A</u>dipocyte-specific <u>D</u>ouble ATGL and HSL <u>K</u>nock-<u>O</u>ut mice (ADKO) were generated by intercrossing *Pnpla2*^flx/flx^::*Adiponectin*-Cre^+/-^ mice with *Lipe*^flx/flx^ mice to obtain *Pnpla2*^flx/flx^*Lipe*^flx/flx^ :: *Adiponectin*-Cre^+/-^ mice. *Pnpla2*^flx/flx^*Lipe*^flx/flx^ littermates were used as controls.

The expression of Cre recombinase, fused to the mutant form of the estrogen receptor (ER) ligand binding domain, and thus gene deletion was induced by oral gavage of 2 mg tamoxifen (Sigma) solubilized in corn oil (Sigma) for five consecutive days. Both CreER^-/-^ and CreER^+/-^ mice obtained tamoxifen treatment at the age of at least 8 weeks. As indicated in the figure legends, studies were performed 6-11 weeks after the first tamoxifen administration. For cold adaptation, gene deletion was induced prior to cold exposure at 5 °C. To maintain gene deletion during cold adaptation, 3 mg tamoxifen per mouse were administered twice per week or once a month, respectively. All mouse models were backcrossed onto a C57Bl/6J background for >10 generations.

### Ethics statement

All animal protocols were performed in accordance with the European Directive 2010/63/EU, the guidelines of the ethics committee of the University of Graz, and were approved by the Austrian Federal Ministry for Science, Research, and Economy (BMWF-66.007/0029-V/3b/2019). Experimental design and reporting for mouse studies follow the ARRIVE guidelines ^54^.

### Mouse husbandry

Mice were bred and maintained under specific pathogen free (SPF) conditions and fed a regular chow diet (R/M-H extrudate, Ssniff). The health status of mice was monitored according to the guidelines of the Federation of European Laboratory Animal Science Associations (FELASA) three times per year using the dirty bedding sentinel program. Regular housing temperatures were maintained between 22-23 °C with a 14 h light / 10 h dark cycle. If possible, mice were maintained in groups of 2-4 mice per cage during maintenance and had *ad libitum* access to water and regular chow diet. After completion of animal protocols, blood was collected via the retro-orbital sinus, mice were euthanized by cervical dislocation, tissues were dissected and immediately processed or flash-frozen in liquid N_2_. Plasma and tissue samples were stored at -80 °C until further analyses.

For all studies, age-matched male littermates were used as controls. Genotype, sex, age, time upon tamoxifen treatment, and number of mice are indicated for each experiment in the appropriate figure legends. Mice were allocated to experimental groups based on their genotype and were treatment-naïve at the time of the study, except for inducible mutant mice receiving tamoxifen treatment.

### Studies at cold

For cold adaptation studies, mice were single-housed and transferred from 22-23 °C to 5 °C for ≥3 weeks without any temperature gradient. Mice had free access to food and water, except otherwise stated.

### Analyses of body temperature

Body temperature was assessed in conscious mice using a rectal probe RET-3 (Physitemp). Core body temperature was studied using implantable telemetry devices (TA-F10, DSI). Telemetry devices were surgically implanted in anesthetized mice (95 µg ketamine and 9.5 µg xylazine per g body weight). Upon surgery, mice were kept on heating plates at 37 °C until they regained consciousness, and were single-housed post-surgery to avoid interference during measurements. Animals received 0.1 mg enrofloxacin and 2 mg ibuprofen per ml drinking water for 3-4 days. Together, mice were allowed to recover from surgery for 7-10 days. Core body temperature was continuously assessed at an interval of 2 min for a period as indicated. Data acquisition and analyses were performed using Dataquest™ A.R.T.™ 4.31.

### Metabolic phenotyping

Oxygen consumption, carbon dioxide production, respiratory exchange ratio (RER), energy expenditure, and locomotor activity were analyzed at given ambient temperatures in unrestrained mice using a laboratory animal monitoring system supplemented with an ActiMot module (PhenoMaster). Mice were familiarized to single housing and drinking flasks for ≥24 h before metabolic phenotyping. Thereafter, the experiments were performed for four consecutive days. Data of the first recorded 12 h were excluded from all analyses. The data were separated on the basis of regular light and dark cycle, averaged or summed up (locomotor activity) of indicated time periods, and analyzed according to guidelines^55^. Food intake was determined manually by weighing the food from single-housed mice every day at 9 AM and 6 PM representing dark and light periods, respectively.

### Analyses of recruitable adrenergic thermogenesis

To test recruitable adrenergic thermogenesis as the key read-out for UCP1-dependent non-shivering thermogenesis (NST)^27^, the metabolic rate after administration of the β_3_-adrenoreceptor agonist CL316,243 (CL) was monitored using PhenoMaster. Mice were anesthetized with pentobarbital (75 mg per kg body weight, *i*.*p*. injection), which does not interfere with UCP1-dependent NST^56,57^. The basal metabolic rate was monitored for subsequent 30 min until reaching steady-state. Then, CL was injected *i*.*p*. using a dosage of 1 mg CL per kg body weight. The subsequent increase in metabolic rate was monitored for 60-80 min. Metabolic rates were measured at an ambient temperature of ∼28-30 °C in order not to underestimate thermogenesis^57^. Control and knockout mice were analyzed in parallel in an interval of 2 min (1 min per cage). Data were analyzed using PhenoMaster software and are expressed as VO_2_ (ml/h).

### Metabolic tracer studies

To trace glucose uptake, 10 µCi of 2-[1,2-^3^H (N)]-deoxy-D-glucose (NEN Radiochemicals) were *i.v.* injected to *ad libitum* fed mice. To trace FAs, [1-^14^C]-*R*-2-Br-palmitic acid (American Radiochemicals) was complexed in a saline solution containing 1% BSA. Radioactively labelled TG rich lipoproteins (TRL) particles were prepared as described^58,59^ with minor modifications. Briefly, TRL-like particles were prepared from 34.2 mg of total lipid including 25.2 mg triolein, 8.172 mg phosphatidylcholine, 0.828 mg lysophosphatidylcholine supplemented with 36 μCi ^3^H-triolein (NEN Radiochemicals). The lipids were dried and sonicated in 3.6 ml of phosphate-buffered saline (Virsonic 475, Virtis) to produce TRL-like particles. Prior to FA and TRL tracer experiments, mice were fasted for 2 h in cold. Then, mice were *i.v*. injected via the retroorbital sinus with 100 µl saline solution containing 1 µCi *R*-2-Br-palmitic acid or 2 µCi ^3^H-TRL particles. After 15 min of tracer administration and in cold, blood was collected, mice were euthanized, and tissues were dissected and stored at -20 °C until further analyses. To determine the radioactivity, an aliquot of total tissue was weighed and lysed overnight in 0.5-1 ml Solvable™ based on tissue weight at 55 °C with constant shaking. Aliquots of 200 µl of tissue lysates were added to 5 ml of scintillation cocktail and the radioactivity was analyzed using liquid scintillation counting (HIDEX 600 SL). The radioactivity was calculated per total tissue.

### *In vivo* lipolysis

To assess whole body lipolysis, blood was collected in the *ad libitum* fed (basal) state and 15 min post-CL injection (*i.p.*). As a readout for lipolysis, plasma FA (NEFA-HR, Fujifilm Wako) and plasma glycerol (free glycerol reagent, Sigma) concentrations were quantified using enzymatic colorimetric methods following manufacturer‘s instructions.

### Histological analyses

Fat tissues were fixed in 4% buffered formaldehyde and embedded in paraffin. Prior to hematotoxylin/eosin (H/E) or immunohistochemistry (IHC), 2 µm thick sections were deparaffinized using xylene and rehydrated using descending dilutions of ethanol and distilled water. For H/E staining, the sections were then stained with hematoxylin. After washing, the sections were stained with eosin-phloxin, dehydrated with ethanol, cleared using butylacetate, mounted, and visualized. For IHC, the sections were placed in EDTA Na buffer (0.5 M/pH: 8) and heated for 40 min at 150 W in a microwave for antigen retrieval. After cooling, the sections were washed using distilled water and phosphate-buffered saline, and blocked using 3% H_2_O_2_ in methanol for 15 min at RT to block endogenous peroxidase activity. Thereafter, the slides were incubated with primary antibodies against UCP1 (1:100 dilution in Dako antibody diluent) for 60 min at RT. The slides were then incubated with EnVision™ Dual Link System-HRP (#K5007, DAKO, Agilent Technologies) for 30 min at RT and visualized using the AEC Substrate Chromogen (#958D-20, Cell Marque). After washing, the slides were counter-stained with hematotoxylin to stain nuclei, mounted with aquatex (Merck), and visualized.

### Transmission electron microscopy

Mice were anesthetized using 258 mg ketamine and 25.8 mg xylazine per g body weight. After ensuring deep anesthesia, mice were perfused with ice-cold phosphate-buffered saline, and subsequently with 4% paraformaldehyde. Then, BAT was dissected, cut into small fragments, and fixed in 2.5% (w/v) glutaraldehyde and 2% (w/v) paraformaldehyde in 0.1 M cacodylate buffer at pH: 7.4 for 3 h, and post-fixed in 2% (w/v) osmium tetroxide for 3 h at RT. After dehydration in graded series of ethanol, the tissues were infiltrated (ethanol and TAAB epoxy resin, pure TAAB epoxy resin) and placed in TAAB epoxy resin for 8 h, transferred into embedding moulds, and polymerized for 48 h at 60 °C. Ultrathin sections of 70 nm were cut with a UC 7 Ultramicrotome (Leica Microsystems) and stained with lead citrate for 5 min and platin blue for 15 min. Electron micrographs were taken using a Tecnai G2 transmission electron microscope (FEI) with a Gatan ultrascan 1000 charge coupled device (CCD) camera (-20 °C; acquisition software Digital Micrograph). Acceleration voltage was 120 kV. Large areas of BAT at high resolution were taken with the scanning transmission electron microscopy mode (STEM) of a field emission scanning electron microscope (ZEISS FE-SEM Sigma 500) in combination with ATLAS TM.

### Analyses of gene expression and relative mitochondrial content using RT-qPCR

Total RNA was extracted from snap-frozen tissues using TRIzol™ reagent following the manufacturer‘s instructions. The quality of RNA was determined by standard agarose gel electrophoresis. RNA concentrations were analyzed using NanoDrop® microvolume spectrophotometer (Thermo Fisher Scientific). To avoid DNA contaminations, 1 µg RNA was digested with 1U Dnase I at 37 °C for 10 min followed by addition of 1 µl 50 mM EDTA and heat inactivation of the enzyme at 75 °C for 10 min. Thereafter, 1 µg RNA was reverse transcribed into single-stranded cDNA using LunaScript RT SuperMix Kit (NEB). For gene expression or relative mitochondrial content, 8-40 ng cDNA, 10 pmol of forward and reverse primers, SYBR Green (Thermo Fisher Scientific), and StepOnePlus™ Real-Time PCR System (Thermo Fisher Scientific) were used for the PCR reaction. Relative gene expression and mitochondrial content were analyzed using the 2^ΔΔ^-Ct method. Relative gene expression was normalized to *Tbp, 36b4, or AdipoQ* (to account for adipocyte-specific gene expression). Relative mtDNA content was determined as ratio of the copy numbers from the mtDNA encoded gene (MtCO1) to the nuclear DNA encoded gene (Ndufv1). A list of gene-specific primers is available in Extended Table 1.

### Fractionation of BAT (adipocytes, stromavascular fraction) and DNA isolation

Freshly dissected BAT depots were cleared of WAT. Whole BAT depots were cut into small pieces and digested as previously described^20^ using collagenase 2 (C2-22, Sigma) and dispase 2 (Sigma). Subsequently, DNA was isolated from brown adipocytes and the stromavascular fraction (SVF) using the DNeasy blood & tissue kit (Qiagen). DNA concentration was determined using a NanoDrop spectrophotometer.

### Isolation of different cellular fractions from BAT (mitochondria/lysosomes, plasma membrane)

For the enrichment of mitochondria, BAT depots were cut into small pieces and homogenized in ice-cold solution A (0.25 M sucrose, 1 mM EDTA, 1 mM dithiothreitol, pH: 7.0, supplemented with 20 µg/ml leupeptin, 2 µg/ml antipain, 1 µg/ml pepstatin), filtered through a 100-µm cell strainer, and centrifuged at 1,000 x *g* for 10 min and 4 °C. Pellets containing nuclei and unbroken cells were discarded. The supernatant excluding floating fat was collected and centrifuged at 12,000 x *g* for 30 min and 4 °C. This supernatant was discarded and pellets containing mitochondria were washed twice in ice-cold solution A and centrifuged at 12,000 x *g* for 15 min and 4 °C. The pellets were resuspended in solution A and the protein content was analyzed using Protein Assay dye and BSA as standard.

For the isolation of the plasma membranes, BAT depots were cut into small pieces and homogenized in an ice-cold Tris-sucrose buffer (50 mM Tris-Cl, 250 mM sucrose, pH: 7.4). The cut tissue pieces were homogenized in a pre-cooled Dounce homogenizer using 15 strokes. The collected tissue homogenates were filtered through a 100-µm cell strainer and centrifuged at 800 x *g* for 10 min and 4 °C. The pellet, containing nuclei and unbroken cells, was discarded. The supernatant was collected and centrifuged at 16,000 x *g* for 30 min and 4 °C to pellet mitochondria. Then, the resulting supernatant excluding the floating fat and the mitochondria pellet was centrifuged at 100,000 x *g* (Beckman Optima^TM^ TLX ultracentrifuge) for 2 h at 4 °C. The supernatant containing the cytosolic fraction was collected and the pellet containing the plasma membrane fraction was resuspended in Tris-sucrose buffer by sonicating in a water bath for 15 min. The resuspended pellets were re-centrifuged at 100,000 x *g* for 45 min at 4 °C. The pellets were resuspended in Tris-sucrose buffer for 15 min by sonication in a water bath, followed by several passes through a 26-gauge needle. The protein content was analyzed using Protein Assay dye and BSA as standard.

### Immunoblotting

For BAT and ingWAT, one depot was weighed and homogenized in ice-cold solution A using an Ultra-Turrax® Homogenizer (IKA). One tissue depot was lysed in the respective volume to adjust to a final volume of 1 ml taking into account the volume changes due to the tissue mass increase. The homogenates were filtered through a 100 µm cell strainer to get rid of connective tissue. For delipidation, the homogenates were processed according to Wessel and Flügge^60^ with minor modifications. In brief, 40 µl of tissue homogenate was mixed with 160 µl water. Then, 800 µl of methanol and 200 µl chloroform were added to the homogenate and vortexed vigorously. To facilitate phase separation, an additional 600 µl water was added, and the samples were vortexed vigorously and centrifuged at 12,500 x *g* for 5 min and RT. The upper aqueous phase was removed and 600 µl methanol was added to the remaining lower phase. The samples were again centrifuged, the supernatant was removed, and the protein pellet was air-dried, lysed in 80 µl of SDS sample buffer, and denatured. Tissue homogenates equivalent to 0.25% of total tissue were loaded onto an SDS polyacrylamide gel.

Alternatively, adipose depots were homogenized in ice-cold RIPA lysis buffer (150 mM NaCl, 10 mM Tris-HCl pH: 7.4, 0.1% SDS, 1% Triton X-100, 1% sodium deoxycholate, and 5 mM EDTA pH: 8.0, supplemented with 20 µg/ml leupeptin, 2 µg/ml antipain, 1 µg/ml pepstatin, and phosphatase inhibitor cocktail (PhosSTOP, Roche) using an Ultra-Turrax® and filtered through a 100-µm cell strainer. The samples were centrifuged at 12,000 x g for 30 min and 4 °C, the infranatant excluding the fat layer was collected, and the protein content was determined using a BCA protein assay and BSA as standard. Protein samples of 1-20 µg were solubilized, denatured in SDS sample buffer, and loaded onto an SDS polyacrylamide gel.

Proteins were resolved by SDS-PAGE using a 10 or 12.5% SDS polyacrylamide gel, and transferred onto a polyvinylidene fluoride (PVDF, 0.45 µm) transfer membrane in CAPS buffer (10 mM CAPS, 10% methanol, pH: 11.0). The membranes were blocked with blocking buffer using 10% blotting grade milk powder (Roth) in TST (50 mM Tris-HCl, 0.15 M NaCl, 0.1% Tween-20, pH: 7.4) for ≥1 h at RT or overnight at 4 °C. Antibodies are listed in Extended Table 2. Protein expression was visualized by enhanced chemiluminescence using Clarity™ Western ECL Substrate and ChemiDoc™ Touch Imaging System (Bio-Rad). Signal intensities were quantified by densitometric analyses using Image Lab software 5.2.14. Total protein stain was performed using Coomassie stain solution for a few seconds and subsequent washing steps in de-staining solution.

### *In vitro* enzyme activity assays

Low-fat tissue homogenates were prepared in solution A and centrifuged at 20,000 x *g* for 30 min at 4 °C to obtain low-fat tissue infranatants. TG hydrolase activities were assessed as described^61^. The TG substrate consisted of 1.67 mM triolein and 188 µM phosphatidylcholine:phosphatidylinositol (3:1; Sigma) supplemented with ∼1,5 x 10^3^ cpm/nmol triolein [9,10-^3^H(N)] per reaction. The lipids were mixed, dried under a N_2_ stream, and emulsified by sonication (Virsonic 475) on ice in 100 mM potassium phosphate buffer at pH: 7.0. The substrates were then adjusted to 5% FA-free BSA (A6003, Sigma) and kept on ice until usage. Lipolytic activities were inhibited using small-molecule inhibitors specific for ATGL (Atglistatin®, ATGLi; final concentration: 40 µM^62^), HSL (HSLi; final concentration: 10 µM; NovoNordisk^13^), or using the non-specific serine hydrolase inhibitor orlistat (final concentration: 10 µM, Sigma). Control incubations contained DMSO.

To determine neutral TG hydrolase activities, 25 µl of the substrate were mixed with 20-50 µg sample to obtain a final volume of 50 µl. As blank, 25 µl solution A were incubated under the same conditions as the samples. The reaction mixtures were incubated in a water bath at 37 °C for 1 h. Then, 650 µl of methanol/chloroform/heptane (10/9/7, v/v/v) and 200 µl of 0.1 M potassium-carbonate/0.1 M boric acid (pH: 10.5) were added to terminate the reaction. The reaction mixture was intensively vortexed and centrifuged for 10 min at 800 x *g*. An aliquot of 200 µl of the upper aqueous phase was collected in 2 ml scintillation cocktail and the radioactivity was analyzed using liquid scintillation counting (Tri-Carb 2100TR). Corrections for background were accounted using counts obtained from blank incubation. The samples from control and mutant mice were run in parallel and analyzed in triplicate or quadruplet. TG hydrolase activities were expressed as released FA per h and normalized to mg tissue protein.

### Analyses of oxygen consumption rates

Freshly dissected BAT (cleard from WAT) and ingWAT depots were collected, minced into small pieces, and homogenized in ice-cold 250 mM sucrose solution in a total volume of 1 ml for 2-3 s using an Ultra-Turrax® Homogenizer. Prior to the analyses, the homogenates were freed from connective tissue using a 100-µm cell strainer. Oxygen consumption rates were measured at 37 °C using polarographic oxygen sensors in a two-chamber Oxygraph (OROBOROS^®^ Instruments). A tissue volume equivalent of 20-100 µl (total volume = 1 ml per ½ tissue depot) was incubated in respiration buffer containing 125 mM sucrose, 20 mM potassium TES buffer (pH: 7.2), 2 mM MgCl_2_, 1 mM EDTA, 4 mM KH_2_PO_4,_ 3 mM malate, and 0.1% FA-free BSA. Respiration was analyzed by the following protocol by adding substrates sequentially upon reaching steady-state: 5 mM glycerol-3-phosphate (G3P; complex-II) and 10 µM cytochrome C, and 2 mM guanosine 5-diphosphate (GDP, dissolved in 20 mM TES buffer pH: 7.2), and 2.5 µM antimycin A (Ama). Substrates and inhibitors were prepared according to Oroboros protocols. Data acquisition and analyses were performed using DatLab 5.1.1.91. Analyses of control and mutant mice were prepared and run in parallel. Tissue respiration rates using tissue homogenates were calculated for the entire depot per mouse and were expressed as nmol/min/entire depot.

### Quantification and statistical analyses

Figures were prepared using GraphPad Prism 10.5.0. All data are shown as means + S.D. except otherwise stated. In each experiment, *n* defines the number of mice. Statistically significant differences between two groups were determined by unpaired, two-tailed Student’s *t*-test. Simple linear regression analyses were used to assess differences between slope elevations. Group differences were considered statistically different for *, *p*<0.05, **, *p*<0.01, and ***, *p*<0.001. Statistical tests, exact values of *n*, definitions of center, dispersion, and significance are indicated in the figure legends.

## Acknowledgements

We thank Sabrina Hütter and Silvia Schauer for excellent technical assistance as well as Dr. Kathrin A. Zierler and her team for excellent animal care and genotyping. This work was supported by the PhD training program doc.fund „Molecular Metabolism-MOBILES“ 10.55776/DOC50 (R. Zimmermann, R. Zechner), 10.55776/PAT3403323 (U.T.), the SFB „Lipid Hydrolysis“ 10.55776/F73 (D.Kr., R.Zi./U.T., R.Ze./R.S.), and the excellence cluster 10.55776/COE14 (R.S., M.S.) – all funded by the Austrian Science Fund FWF, the Leducq Foundation Grant 12CVD04, and the Louis Jeantet Foundation. For the purpose of open access, the authors have applied a CC BY public copyright license to any author accepted manuscript version arising from this submission. We thank the University of Graz for financial support and gratefully acknowledge the support of NAWI Graz, BioTechMed-Graz, and the *Field of Excellence* BioHealth-University of Graz, Graz, Austria.

## Author contributions

R.S. and R.Ze. conceptualized the study. Y.M. and R.S. conducted most of the experiments and analyzed the data. A.P., P.H., and W.K. performed immunoblotting and activity assays. D.Ko. and G.H. analyzed electron microscopy and histological analyses, respectively. N.V. and D.Kr. provided indirect calorimetry devices. R.S. wrote the manuscript, Y.M., N.V., M.S., U.T., R.Zi., D. Kr., R.Ze., and R.S. reviewed and edited the manuscript.

---

**Extended Data Fig. 1 – related to Fig. 2.**
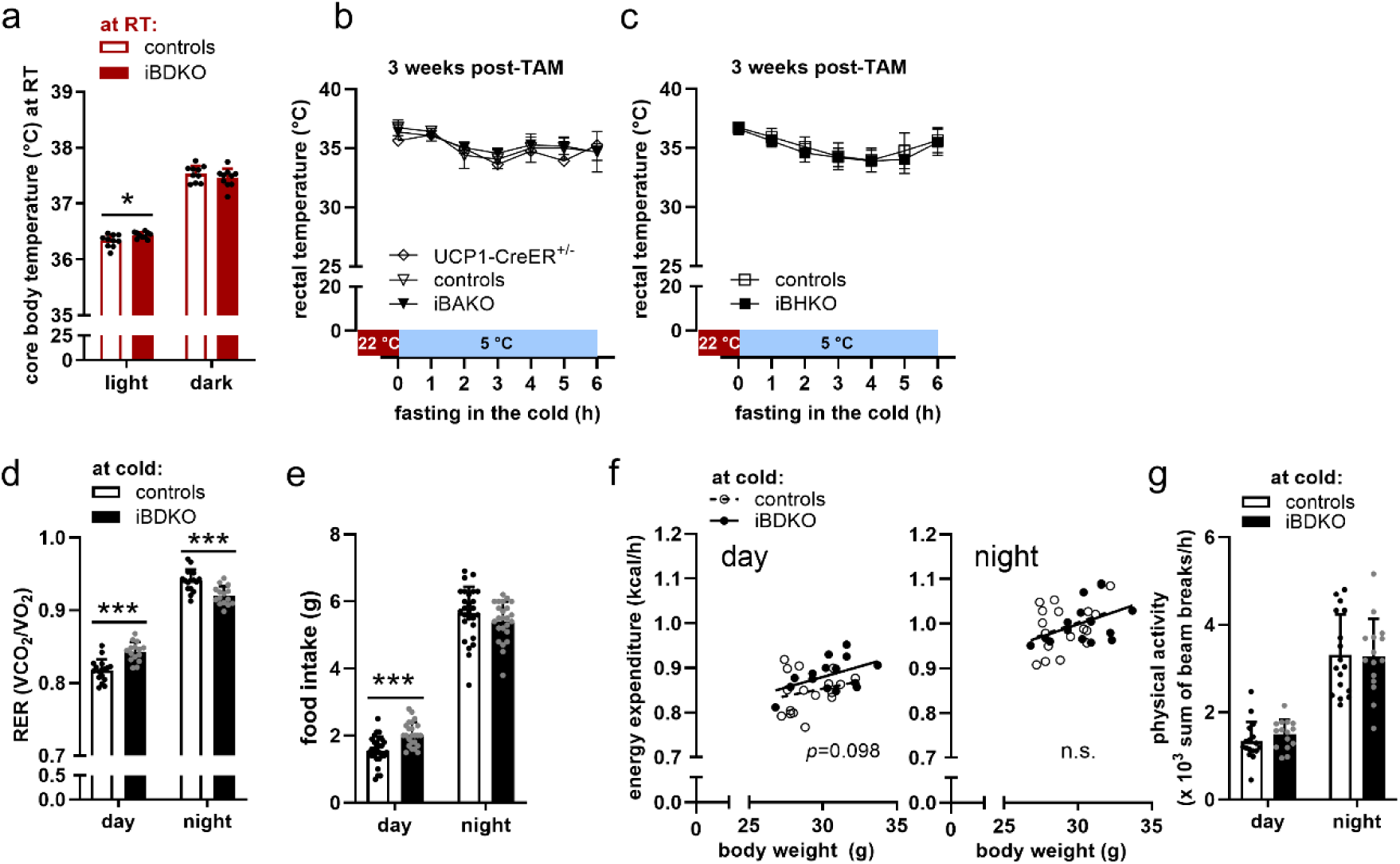
Metabolic analyses of iBDKO mice. (**a**) Core body temperature at room temperature (RT) (n = 10), (**b**) Body temperature from tamoxifen-inducible <u>B</u>rown-adipocyte specific <u>A</u>TGL <u>K</u>nock-<u>O</u>ut mice (iBAKO, n = 3-4) and *Ucp1*-CreER^+/-^ (n = 3) mice 3 weeks upon gene deletion using tamoxifen (post-TAM). Mice were previously housed at room temperature at 22 °C (red area) and acutely shifted to 5 °C during fasting (light blue area). (**c**) Body temperature from tamoxifen-inducible <u>B</u>rown-adipocyte specific <u>H</u>SL <u>K</u>nock-<u>O</u>ut mice (iBHKO, left graph, n = 8-13) mice 3 weeks upon gene deletion using tamoxifen (post-TAM). Mice were previously housed at room temperature at 22 °C (light red area) and acutely shifted to 5 °C during fasting (light blue area). (**d**) Respiratory exchange ratio (RER). Non-shaded and grey shaded areas represent day and night, respectively. (n = 15-17). (**e**) Food intake (n = 24-27). (**f**) Energy expenditure at day (left) and night (right) (n = 15-17). n.s., not significant. (**g**) Physical activity (n = 15-17). For the analyses of core body temperature, mice were implanted with telemetry transmitters. Body temperature was assessed using a rectal probe. iBDKO male mice aged 20-24 weeks and 6-10 weeks post-TAM were exposed to 5 °C for ≥3-weeks. iBAKO males were aged ≥45 weeks and iBHKO males were aged 13-15 weeks at indicated times post-TAM. Data are means + SD (except for time-course in B showing mean only). Statistical analyses were performed using Student’s *t*-test. Simple linear regression analyses were used to assess differences between slope elevations (D). *, *p*<0.05 and ***, *p*<0.001.

**Extended Data Fig. 2 – related to Fig. 3.**
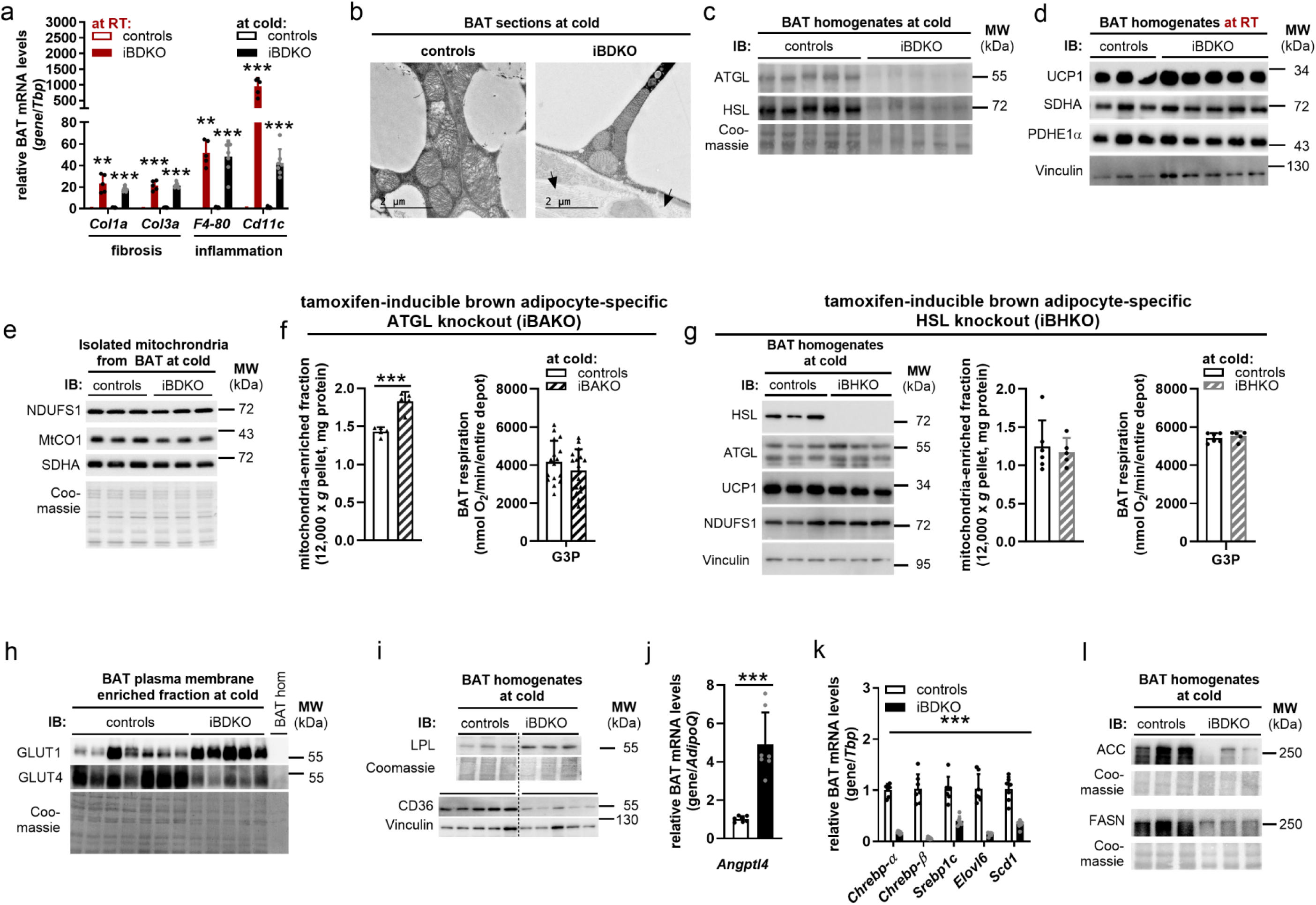
iBDKO mice have impaired BAT thermogenic capacity upon cold adaptation. (**a**) Gene expression in BAT from controls and iBDKO mice housed at room temperature (RT, n = 3-5) and during cold adaptation (cold, n = 7). (**b**) Transmission electron microscopy of BAT from cold adapted controls and iBDKO mice. Arrows indicate collagen depositions. Bar graph = 2 µm. (**c**, **d**) Immunoblot (IB) of BAT homogenates from mice upon cold adaptation (cold) and room temperature (RT). Delipidated volume equivalents of 0.25% of total BAT were loaded per lane. (**e**) Immunoblot (IB) from isolated mitochondria from BAT upon cold adaptation. Per lane, 2.5 µg protein was loaded. (**f**, **g**) Protein content in the mitochondria-enriched fraction (12,000 x *g* pellet) isolated from BAT from mice upon cold adaptation and respiration using glycerol-3-P (G3P) as substrate in BAT homogenates from mice upon cold adaptation from (**f**) tamoxifen-inducible <u>B</u>rown-adipocyte specific <u>A</u>TGL Knock-Out mice (iBAKO, n = 5/mitochondria; n = 16/respiration) and (**g**) tamoxifen-inducible <u>B</u>rown-adipocyte specific <u>H</u>SL <u>K</u>nock-<u>O</u>ut mice (iBHKO, n = 5-6). Immunoblot (IB) of BAT homogenates from iBHKO mice upon cold adaptation (cold). (**h**) Immunoblot (IB) of plasma membrane fractions isolated from BAT. Per lane, 5 µg protein was loaded. (**i**) Immunoblot (IB) of BAT homogenates upon cold adaptation. Volume equivalents of 0.25% of total BAT were loaded per lane. (**j**, **k**) Gene expression in BAT during cold adaptation (n = 7). (**l**) Immunoblot (IB) of BAT homogenates. Delipidated volume equivalents of 0.25% of total BAT were loaded per lane. RT-housed male mice were 21-24 weeks of age and 7 weeks post-TAM. Cold-adapted male mice were 16-23 weeks of age and 6-11 weeks post-TAM that were cold-adapted at 5 °C for ≥3 weeks. Data are means + SD. Statistical analyses were performed using Student’s *t*-test. **, *p*<0.01 and ***, *p*<0.001.

**Extended Data Fig. 3 – related to Fig. 4.**
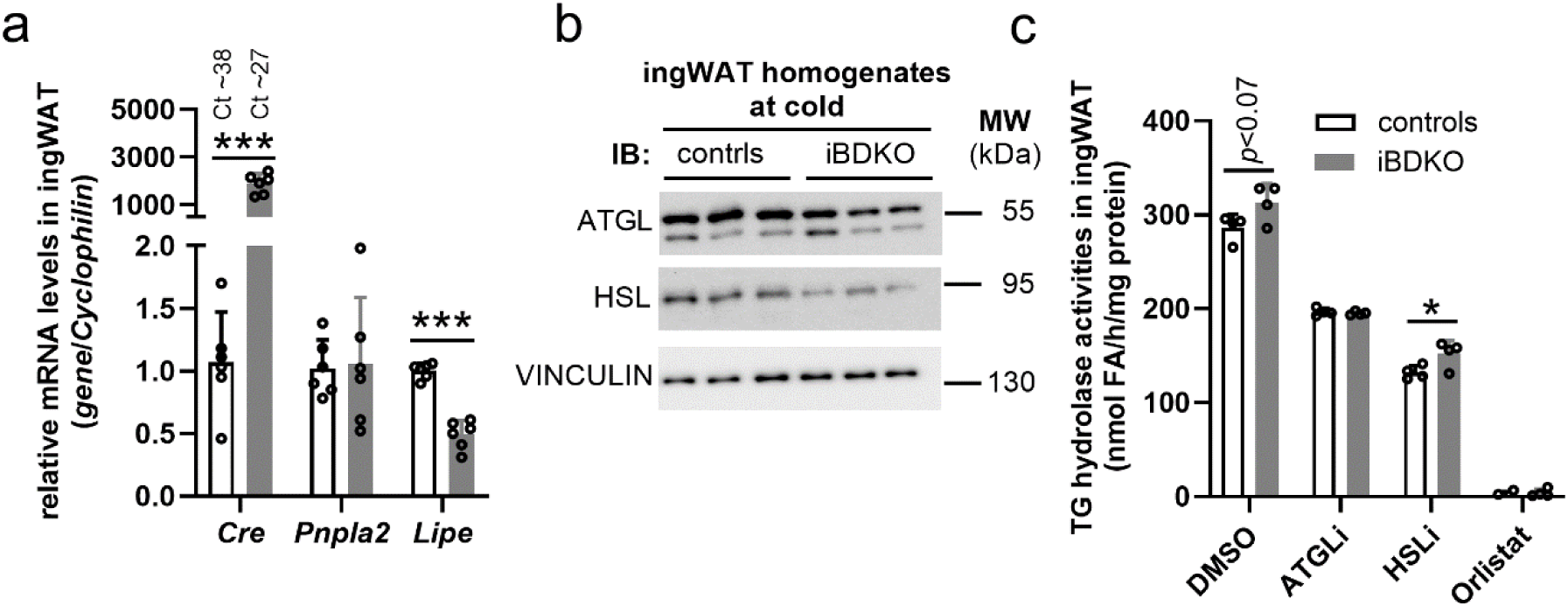
Browning of inguinal WAT in cold-adapted iBDKO mice. (**a**) Gene expression in ingWAT (n = 6). (**b**) Immunoblot (IB) of ingWAT homogenates. (**c**) TG hydrolase activities were determined in a pool of fat-free infranatants from of ingWAT homogenates (n = 5-6) using a radioactive labeled triolein substrate emulsified with phospholipids. Enzymatic activities were determined in the absence or presence of inhibitors specific for ATGL (ATGLi) or HSL (HSLi) or the non-specific serine hydrolase inhibitor orlistat. Male mice of 20-21 weeks of age and 6-8 weeks post-TAM were exposed to 5 °C for ≥3 weeks. Data are means + SD. Statistical analyses were performed using Student’s *t*-test. *, *p*<0.05 and ***, *p*<0.001.

**Extended Data Fig. 4 – related to Fig. 5.**
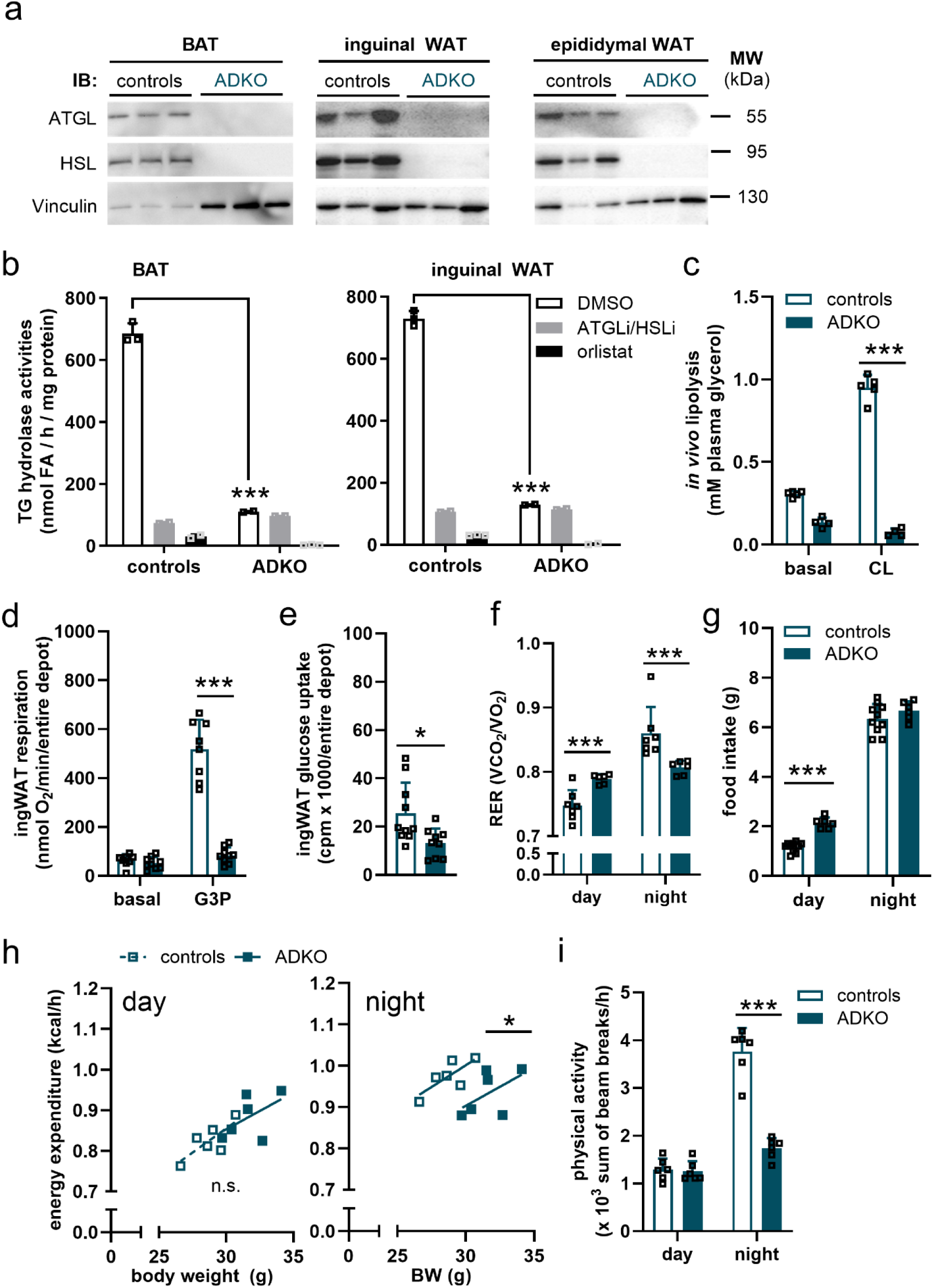
(**a**) Immunoblot (IB) from *ad libitum* fed mice housed at RT. Volume equivalents of 0.25% of total tissues were loaded per lane. (**b**) TG hydrolase activities were determined in a pool of fat-free infranatants from BAT and ingWAT homogenates (n = 4) using a radioactive labeled triolein substrate emulsified with phospholipids. Enzymatic activities were determined in the absence or presence of inhibitors specific for ATGL (ATGLi) or HSL (HSLi) or the non-specific serine hydrolase inhibitor orlistat. Technical replicates are shown. (**c**) *In vivo* lipolysis during unstimulated (basal) and CL 316,243 (CL) stimulated conditions. Blood was collected from *ad libitum* fed mice (n = 5) for basal und 15 min upon *i.p.* injection of 1 mg CL/kg body weight. Plasma glycerol was determined using an enzymatic colorimetric assay. (**d**) Respiration in ingWAT homogenates upon cold adaption (n = 8-9). (**e**) Glucose uptake using ^3^H-2-deoxyglucose in ingWAT upon cold adaption (n = 9-10). (**f**) Respiratory exchange ratio (RER) upon cold adaption (n = 6-7). (**g**) Food intake (n = 6-10). (**h**) Energy expenditure at day (left) and night (right) (n = 6). (**i**) Physical activity (n = 6). Male mice aged 14-20 weeks were used for the study. For cold adaptation, mice were housed at 5 °C for ≥3 weeks. Data are presented as means + SD. Statistical analyses were performed using Students *t*-test. Simple linear regression analyses were used to assess differences between slope elevations (D). *, *p*<0.05 and ***, *p*<0.001.

**Extended Data Table 1.**
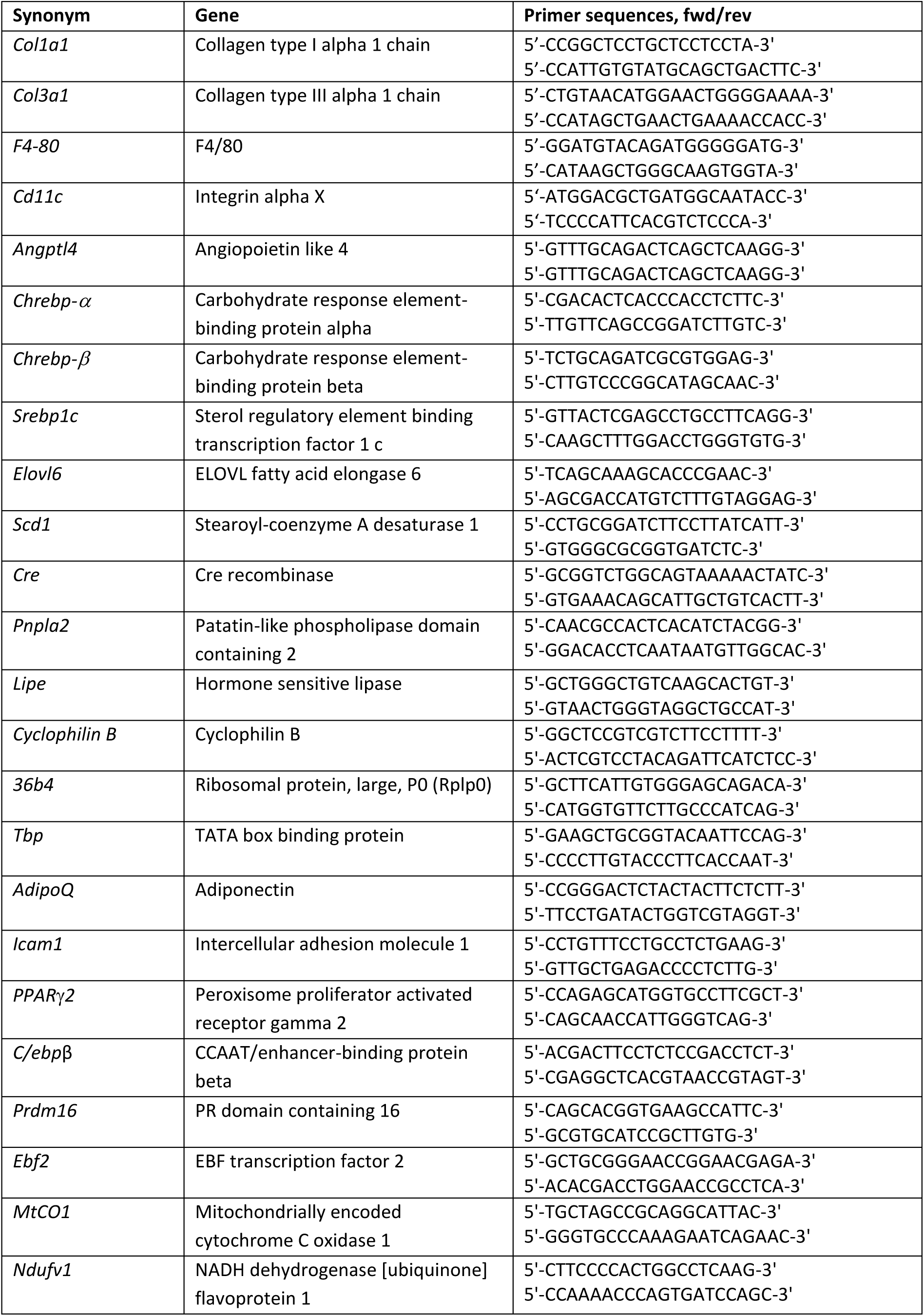
List of RT-qPCR primers.

**Extended Data Table 2.** List of antibodies.

| Antibodies | Source | Identifier |
| --- | --- | --- |
| Adipose triglyceride lipase (ATGL) | Cell Signaling | Cat # 2138;<br>RRID:AB_2167955 |
| Hormone-sensitive lipase (HSL) | Cell Signaling | Cat # 4107;<br>RRID:AB_2296900 |
| Uncoupling protein 1 (UCP-1) | Abcam | Cat # ab10983;<br>RRID:AB_2241462 |
| NADH:ubiquinone oxidoreductase core subunit S1 (NDUFS1) | Abcam | Cat # ab157221;<br>RRID:AB_2857900 |
| Pyruvate dehydrogenase E1 alpha (PDHE1a) | Abcam | Cat # 177461;<br>RRID:AB_2137173 |
| Succinate dehydrogenase complex flavoprotein subunit A (SDHA) | Abcam | Cat # ab14715;<br>RRID:AB_301433 |
| Cytochrome c oxidase subunit 4 (COXIV) | Cell Signaling | Cat #4844;<br>RRID: AB_2085427 |
| Glucose transporter GLUT1 | Cell Signaling | Cat #12939;<br>RRID: AB_2687899 |
| Glucose transporter GLUT4 | Abcam | Cat #ab654;<br>RRID: AB_305554 |
| Lipoprotein lipase (LPL) | Gift from S.G. Young |  |
| CD36 | Abcam | Cat # ab133625;<br>RRID:AB_2716564 |
| Tyrosine hydroxylase (TH) | Millipore | Cat #MAB318;<br>RRID:AB_2201528 |
| Mitochondrially encoded cytochrome C oxidase 1 (MtCO1) | Abcam | Cat# ab14705;<br>RRID :AB_204810 |
| Acetyl-CoA carboxylase (ACC) | Cell Signaling | Cat# 3676;<br>RRID:AB_2219397 |
| Fatty acid synthase (FASN) | Cell Signaling | Cat# 3180;<br>RRID:AB_2100796 |
| Vinculin | Sigma-Aldrich | Cat# V9131;<br>RRID:AB_477629 |
| GAPDH | Cell Signaling | Cat # 2118;<br>RRID:AB_561053 |
| Mouse-HRP | Sigma-Aldrich | Cat # NA931;<br>RRID:AB_772210 |
| Goat-HRP | Millipore | Cat # AP180P;<br>RRID:AB_92573 |
| Rabbit-HRP | Cell Signaling | Cat # 7074;<br>RRID:AB_2099233 |

